# Structural rearrangements of a caspase-like protease TPR-CHAT govern virus-host discrimination during type III-E CRISPR-Caspase immunity

**DOI:** 10.1101/2022.09.03.506347

**Authors:** Ning Cui, Jun-Tao Zhang, Zhuolin Li, Xiao-Yu Liu, Chongyuan Wang, Hongda Huang, Ning Jia

## Abstract

The RNA-targeting type III-E CRISPR-gRAMP effector forms a complex with a caspase-like protease TPR-CHAT, but the mechanistic details of their functional relationship remain unknown. Here, we report on cryo-EM structures of gRAMP^crRNA^ and gRAMP^crRNA^-TPR-CHAT complexes, before and after either self or non-self RNA target binding, elucidating mechanisms underlying RNA-targeting and non-self RNA-induced protease activation. Noteworthy, the associated TPR-CHAT adopts a strikingly distinct conformation on self versus non-self RNA targets, with nucleotides at position −1 and −2 of crRNA serving as a sensor. Only binding of non-self RNA target activates TPR-CHAT protease, leading to the cleavage of Csx30 protein. Furthermore, given that TPR-CHAT structurally resembles eukaryotic separase, our results implicate an ancient mechanism for separase regulation. Our findings should not only facilitate the development of gRAMP-based RNA manipulation tools, but also lead to a mechanistic understanding of the virus-host discrimination process governed by a caspase-like protease during type III-E CRISPR-Caspase immunity.

## Introduction

CRISPR-Cas systems provide prokaryotes with adaptive immunity against invading viruses and plasmids^1,2^. The immunity is acquired by integrating the invading nucleic acid between repeat sequences of the host CRISPR locus. The CRISPR loci are then transcribed into precursor RNA (pre-crRNA) that is processed into mature CRISPR-derived RNAs (crRNA), which assembles with single-and multi-subunit Cas proteins to form crRNA-effector complexes responsible for detection and subsequent degradation of the invading nucleic acid. Depending on their *cas* gene composition, CRISPR-Cas systems are classified into two major classes (1-2) distributed between six types (I–VI)^3^. Class 1 systems (including types I, III and IV) comprise multi-subunit effector complexes, whereas class 2 systems (including type II, V and VI) are composed of single-subunit protein effectors. Of all known CRISPR-Cas systems, RNA-targeting effector complexes have been identified in type III, V and VI systems^4–6^, of which the single-subunit effector type VI Cas13 has been widely used for RNA detection, editing and knockdown^7^. However, upon recognition of the RNA targets, Cas13 exhibits a non-specific RNA cleavage activity resulting in degradation of both viral and host RNA^5,8,9^. The nonspecific, collateral RNA cleavage activity associated with Cas13 have been shown to be toxic to multiple mammalian cell types^9–11^.

Remarkably, a recent characterized type III-E subtype contains a single-protein effector that exhibits specific target RNA cleavage activity with no collateral activity and cell toxicity^3,12,13^, representing a new promising RNA-manipulation tool. Recently, Kato et al. have provided a first glimpse into the architecture of a type III-E CRISPR effector complex *Desulfonema ishimotonii* Cas7-11 bound to a target RNA, elucidating the mechanism for target RNA cleavage^14^. However, given that the type III-E effector complex has been shown to equally cleave both the invasive non-self and the host self RNA targets derived from the host CRISPR array^12^, it still remains unknown how the type III-E effector complex discriminates self and non-self RNA target during antiviral defense.

Type III CRISPR-Cas systems have been divided into six subtypes (type III-A to F), of which type III-A, -D, -E and -F contain Csm proteins, whereas type III-B and -C consists of Cmr proteins^3^. In previously reported canonical type III-A/B systems, the mature crRNA consists of a single spacer that is flanked by 8-nt 5’-repeat tag derived from repeat sequence of the host CRISPR array. The non-complementarity between 3’-flanking sequences of the target RNA and 5’-repeat tag of crRNA specifies an invasive non-self RNA target and triggers the immune response, whereas complementarity between these two sequence elements licenses the RNA target derived from the host CRISPR array and prevents self-targeting^15^. Only binding of the invasive non-self RNA target, but not the host self RNA, allosterically activates the ssDNase and cyclic oligoadenylates (cOA) synthetase activities of the type III signature protein Cas10 to degrading the invading nucleic acids^16–20^. Meanwhile, cleavage of the invasive RNA target by Csm3/Cmr4 subunit switches off Cas10 enzymic activities, thereby preventing the potential damage to the host due to persistent enzyme activities^18,21^.

Intriguingly, lacking the type III signature protein Cas10, the III-E CRISPR loci often contain a gene encoding a caspase-like protease TPR-CHAT that contains a Caspase HetF Associated with TPRs (CHAT) domain, which is a caspase family protease and typically involved in programmed cell death, fused with a tetratricopeptide repeat (TPR) domain^3,12,13,22^ (Fig. 1a). The TPR-CHAT protease has been shown to form a stable complex with *Candidatus* “Scalindua brodae” giant Repeat-Associated Mysterious Protein (gRAMP) in type III-E CRISPR-Cas systems^12^, suggesting a functional relation between CRISPR-Cas systems and caspase-like proteases during type III-E CRISPR-Cas antiviral immunity. The TPR-CHAT protease has been implicated in regulating bacteria death by site-specific cleavage and the subsequent activation of a bacterial gasdermin, which executes cell death for antiphage defense^23^, suggestive of an antiviral role of TPR-CHAT protease in type III-E CRISPR-Cas immunity.

**Fig. 1.**
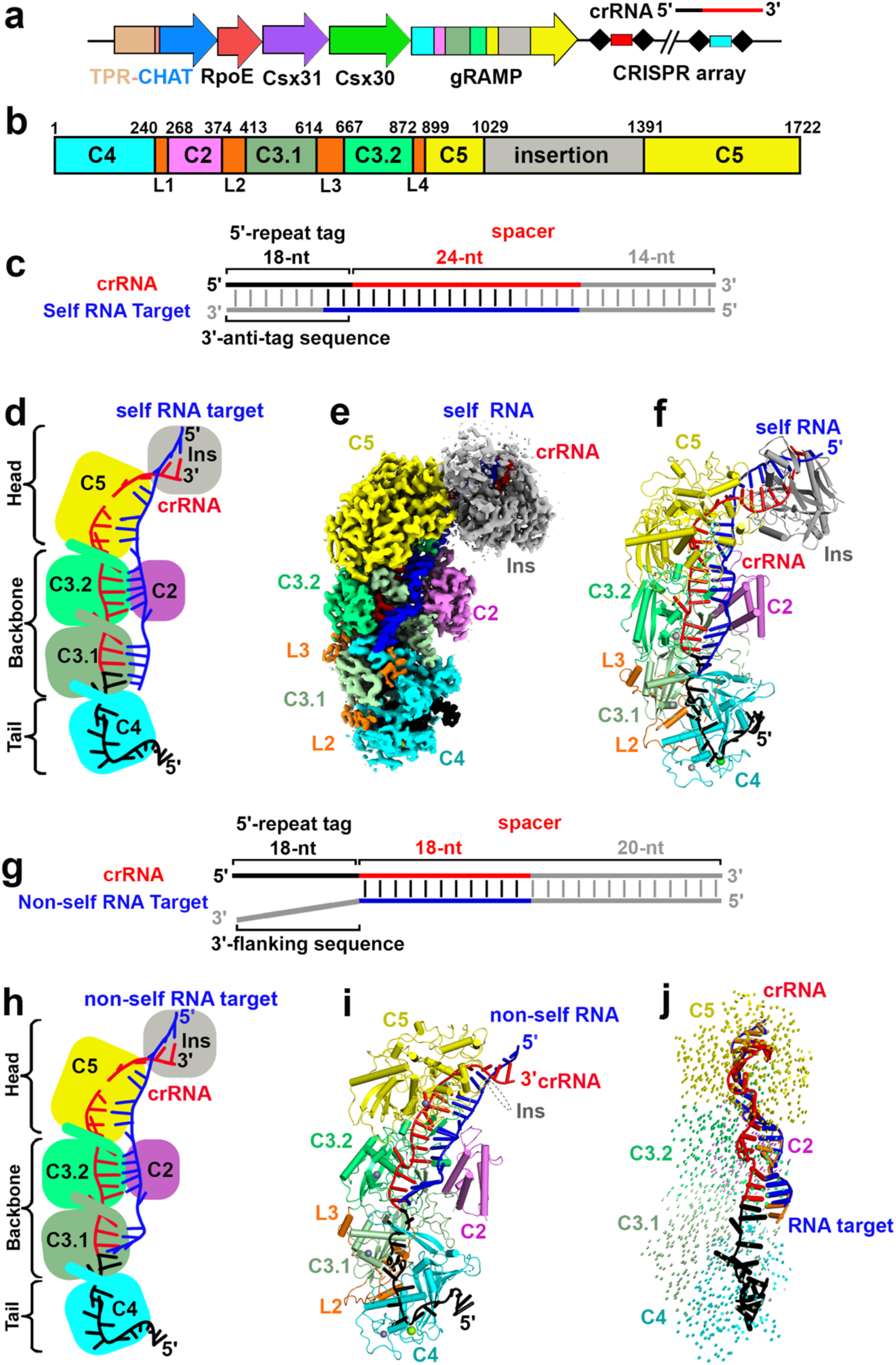
Cryo-EM structures of gRAMP^crRNA^ bound to target RNAs. **a**, The outline of type III-612 E CRISPR-Cas loci from *Candidatus* “Scalindua brodae”. gRAMP and TPR-CHAT genes are 613 colored according to domain organization. The CRISPR locus are composed of the host nucleotide repeats (black diamonds) separated by invading spacer sequences (colored cylinders). **b**, Domain organization of gRAMP protein. C2, Csm2 domain; L1, Loop 1; insertion indicates the insertion domain within Csm5 domain; Other domains are named accordingly. **c**, Schematic representation of pairing between crRNA and self RNA target. Segments that can be traced in gRAMP^crRNA^-target RNA^self^ complex are in color, while disordered segments are in grey. **d, e** and **f**, Schematic (**d**), cryo-EM reconstruction (**e**) and ribbon representation (**f**) of the 2.8 Å structure of gRAMP^crRNA^-target RNA^self^ complex. **g**, Schematic representation of pairing between crRNA and non-self RNA target. Segments that can be traced in gRAMP^crRNA^-target RNA^non-self^ complex are in color, while disordered segments are in grey. **h** and **i**, Schematic (**h**), and ribbon representation (**i**) of the 2.5 Å structure of gRAMP^crRNA^-target RNA^non-self^ complex. **j**, Structural comparison between gRAMP^crRNA^-target RNA^non-self^ and gRAMP^crRNA^-target RNA^self^ complexes. Vector length correlates with the domain movement scale.

However, a mechanistic understanding for the functional relations between type III-E CRISPR-Cas effector complex and the caspase-like protease TPR-CHAT, and also the mechanism for the self versus non-self discrimination in type III-E systems, remain unclear. Here, combining structural biology and biochemistry, we have provided high-resolution snapshots of the type III-E gRAMP^crRNA^ and gRAMP^crRNA^-TPR-CHAT complexes, with or without self or non-self RNA targets bound, thereby revealing mechanisms underlying RNA-targeting and virus-host discrimination governed by the non-self RNA-activated TPR-CHAT protease in type III-E CRISPR-Caspase antiviral immunity.

## Results

### Overall architecture of gRAMP^crRNA^ bound to RNA targets

A previous publication has shown that the type III-E *Candidatus* “Scalindua brodae” gRAMP complex specifically recognizes and cleaves both self and non-self RNA targets^12^. To investigate whether the gRAMP^crRNA^ complex could distinguish self from invasive non-self RNA targets, we first co-expressed the gRAMP gene with a single CRISPR array in *Escherichia coli* cells (Fig. 1a), gRAMP and crRNA formed a stable complex (Extended Data Fig. 1a), and showed that it cleaved both self RNA and non-self RNA targets into two products in a metal dependent manner (Extended Data Fig. 1b). We then reconstructed the complexes of gRAMP^crRNA^ with either self or non-self RNA targets bound by addition of ethylene diamine tetraacetic acid (EDTA) to prevent target cleavage (Extended Data Fig. 1c, d), and determined the cryogenic electron microscopy (cryo-EM) structures of gRAMP^crRNA^-target RNA^self^ and gRAMP^crRNA^-target RNA^non-self^ ternary complexes at 2.8 Å and 2.5 Å, respectively (Fig. 1c-i, Extended Data Fig. 2), with the former containing more observed densities in the head region.

The overall architecture of gRAMP^crRNA^-target RNA ternary complexes adopts a seahorse shape, resembling type III-A CRISPR-Csm complex with fused homologous domains referred to as the head (the Csm5 domain), back-bone (two Cas7-like Csm3 domains and one Cas11-like Csm2 domain), and tail (the Csm4 domain) fused by four Linker domains (L1 to L4), but lacking the type III signature protein Cas10^24–27^ (Fig. 1b, f, i and Extended Data Fig. 3). Densities corresponding to the Csm5 insertion domain in the head region could only be observed in the gRAMP^crRNA^-target RNA^self^ ternary complex, with only parts of the densities being well traced, suggesting its flexibility (Extended Data Fig. 2d). We observed a 36-nt crRNA bound to 18-nt traceable segment of 56-nt non-self RNA target in gRAMP^crRNA^-target RNA^non-self^ ternary complex (Fig. 1g, i). More density was observed for the crRNA-target RNA duplex in the gRAMP^crRNA^-target RNA^self^ ternary complex containing a 42-nt crRNA bound to 24-nt traceable self RNA target (Fig. 1c, f), with the extra observed densities corresponding to the anti-tag sequence that base-pairs with crRNA, and sequences covered by Csm5 insertion domain. Despite the difference between the self and invasive non-self RNA targets, minimal conformational changes of the overall structure are observed between gRAMP^crRNA^-target RNA^non-self^ and gRAMP^crRNA^-target RNA^self^, with a small root-mean-square deviation (r.m.s.d.) of 0.463Å (Fig. 1j), indicating the self versus non-self discrimination could not be manifested by conformational changes of gRAMP^crRNA^ induced by binding of self or non-self RNA targets.

### Assembly of mature crRNA

In the canonical type III systems, the mature crRNA with an 8-nt 5’-repeat tag is produced from pre-crRNA following cleavage by a stand-alone endoribonuclease Cas6^2,28^. Recent publications have shown the Cas7.1 domain in type III-E *D. ishimotonii* Cas7-11 (corresponding to Csm4 domain in gRAMP) is responsible for pre-crRNA processing, leading to the generation of mature crRNA with a 15-nt 5’-repeat tag^13,14^. However, we observed 18-nt 5’-repeat tag in our determined type III-E gRAMP complexes, numbered −18 to −1 according to convention (Fig. 2a), with no cleavage observed between nucleotides −15U and −16C. In addition, the indispensable catalytic residue H43 adjacent to the scissile phosphate at the −15 to −16 step in *D. ishimotonii* Cas7-11 for pre-crRNA processing, is replaced by T45 in gRAMP (Fig. 2b, blue insert), implicating a different pre-crRNA processing mechanism for gRAMP.

**Fig. 2.**
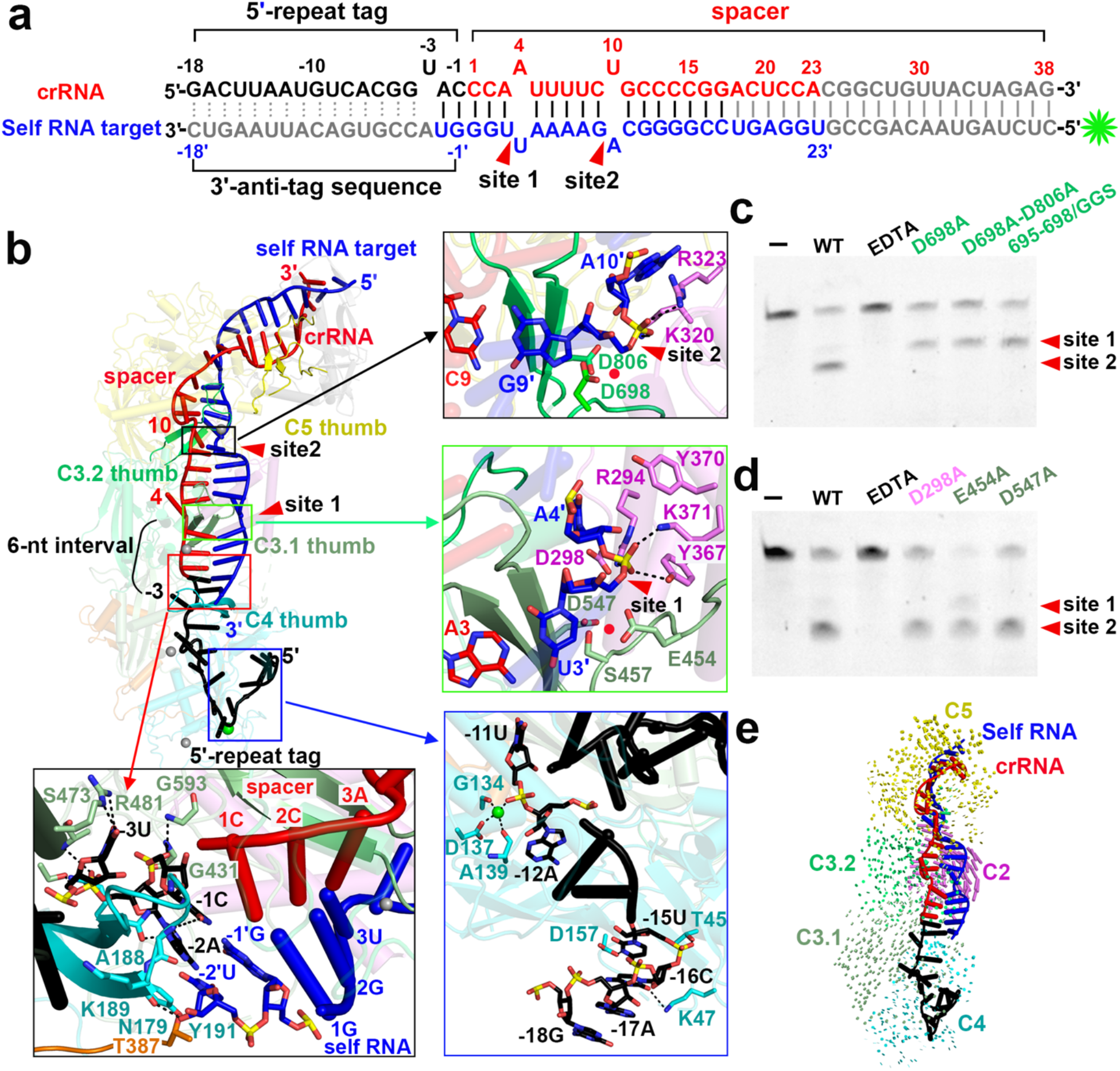
Assembly of crRNA-Target RNA Duplex in gRAMP^crRNA^-Target RNA^self^ complex. **a**, Schematic drawing for the sequences of crRNA and self RNA target. Potential cleavage sites are mapped on the target RNA sequence. Segments that can be traced are in color, while disordered segments are in grey. The green asterisk indicates the fluorescence label at 5’ end of the RNA target for the following target RNA cleavage. **b**, The thumb elements of Csm4 and Csm3 domains kink the crRNA-target RNA duplex every 6th nucleotide, with no kink associated with the thumb of Csm5 domain. The black and blue inserts present interactions that might contribute to RNase activity for the target RNA cleavage at the 9’-10’ step and 3’-4’ step, respectively. The blue insert provides detailed contacts between Csm4 domain and nucleotides at 5’-repeat tag. The green sphere indicates a metal ion. The red insert shows detailed interactions between protein residues and crRNA-target RNA duplex at position −1 and −2. **c** and **d**, Cleavage of target RNA by Csm3.2 (**c**) and Csm3.1 (**d**) mutants in the context of the gRAMP^crRNA^-target RNA complex. The two cleavage sites are indicated as red arrows. **e**, Structure comparison between gRAMP^crRNA^ binary complex and gRAMP^crRNA^ -target RNA^self^ ternary complex based on alignment of the Csm4 domain. Vector length correlates with the domain movement scale.

The nucleotides at position from −1 to −16 in the 18-nt 5’-repeat tag form multiple sequence-specific contacts with residues in Csm4 and Csm3 (Extended Data Fig. 4a), indicating gRAMP recognizes 5’-repeat tag in a sequence dependent manner, which is in accord with the conservation of 5’-repeat tag sequence (Extended Data Fig. 4b). In contrast, a series of sequence-nonspecific contacts are found between crRNA spacer sequence and Csm3.1, Csm3.2 and Csm5 (Extended Data Fig. 4a), accounting for the lack of sequence specificity for spacer sequence recognition. Notably, a metal ion is observed to be coordinated by the phosphates group of −11U, the side chain of D137, and the carboxyl group of G134 and A139 in gRAMP complex (Fig. 2b, blue insert). A previous publication has shown that a magnesium ion critical for guide RNA stabilization is observed in a similar position in prokaryotic Argonautes^29^, suggesting the corresponding metal ion might also function in stabilizing 5’-repeat tag in the gRAMP complex.

### Metal-dependent specific cleavage of target RNA

crRNA guides gRAMP to cleave both self and non-self RNA targets between nucleotides 3’ and 4’ (site 1) and between 9’ and 10’ (site 2) with a 6-nt interval in a metal dependent manner^12^ (Fig. 2a and Extended Data Fig. 1b). The potential position for a divalent cation indispensable for target RNA cleavage is indicated by red ball that sits among the acidic Asp side chains in Csm3.1 and Csm 3.2 domains and the phosphate oxygen (Fig. 2b, black and green inserts). Consistently, mutation of the respective acidic Asp residues (D698 in Csm3.2, and D547 in Csm3.1), which are possibly responsible for coordinating the indispensable metal ion, into alanine respectively, abolished the RNase activity in Csm3.1 and Csm3.2 domains (Fig. 2c, d). Notably, mutation of residue D298 in Csm2 into alanine also abolished RNase activity (Fig. 2d), indicating the critical role of Csm2 for target RNA cleavage. In addition, the two cleavable phosphates are stabilized by residues in Csm2 domain (Fig. 2b, black and green inserts). Notably, these residues undergo significant conformational changes revealed by the structural comparison between the gRAMP^crRNA^-target RNA ternary complex and the gRAMP^crRNA^ binary whose structure is determined at 2.7 Å (Fig. 2e, and Extended Data Fig. 5 and 6a). The mechanism of target RNA cleavage by gRAMP^crRNA^ is reminiscent of previous reported type III multi-subunit effector complexes^30–32^, indicating type III-E gRAMP still retains a similar RNA-targeting mechanism even after domain fusion into a single-protein effector.

Despite the similarity, a kink at position −3 in gRAMP^crRNA^ complex rather than a kink at position −1 in the canonical type III Csm/Cmr complex^24,25,27,32^, separates nucleotides between 5’-repeat tag and spacer sequence of crRNA, acting as a start site for the measurement of 6-nt periodic RNA cleavage sites (Fig. 2a, b). Besides, in canonical type III Csm/Cmr complexes, nucleotides at positions −2 to −5 within 5’-repeat tag form a stacked alignment in a pseudo A-form configuration and are exposed to the solvent, serving as sensors for avoidance of autoimmunity^24–26,32^. But in type III-E gRAMP^crRNA^ complex, nucleotides −1C and −2A within the 5’-repeat tag stack with 1C to 3A within the spacer, which are available for base paring with self RNA target (Fig. 2a and b, red insert), possibly acting as sensors for discriminating self and non-self RNA. However, the gRAMP^crRNA^ complex could equally cleave both self and non-self RNA targets (Extended Data Fig. 1b), with minimal difference observed in RNA catalytic pockets between self and non-self RNA bound gRAMP^crRNA^ complex (Extended Data Fig. 6b), suggesting other factors rather than RNA cleavage is responsible for the self and non-self discrimination in type III-E CRISPR antiviral immunity.

### The caspase-like protease TPR-CHAT binds to the tail of gRAMP^crRNA^ complex

In canonical type III CRISPR-Cas systems, both self and non-self RNA be equally cleaved by Csm3 subunits, but only the invasive non-self RNA could activate the ssDNase and cOA synthetase activities of the signature Cas10 subunit, which is critical for type III antiviral immunity^16–20^. Intriguingly, given that they lack the type III signature *cas10* gene, the type III-E CRISPR locus contains a gene encoding a caspase-like protease TPR-CHAT (Fig. 1a), which forms a stable complex with gRAMP^crRNA^ (Extended Data Fig. 7a). To gain structural insights into how TPR-CHAT protease interacts with the gRAMP^crRNA^ complex, we determined the cryo-EM structure of gRAMP^crRNA^-TPR-CHAT complex at 2.6 Å resolution (Fig. 3a, b, and Extended Data Fig. 7b-d). TPR-CHAT contains an N-terminal TPR domain and a caspase-like protease CHAT domain connected by a Linker domain, and binds the tail of gRAMP^crRNA^ complex mainly through Loop2-mediated electrostatic contacts with a buried interface area of ~1800 Å^2^ (Fig. 3c and Extended Data Fig. 7e).

**Fig. 3.**
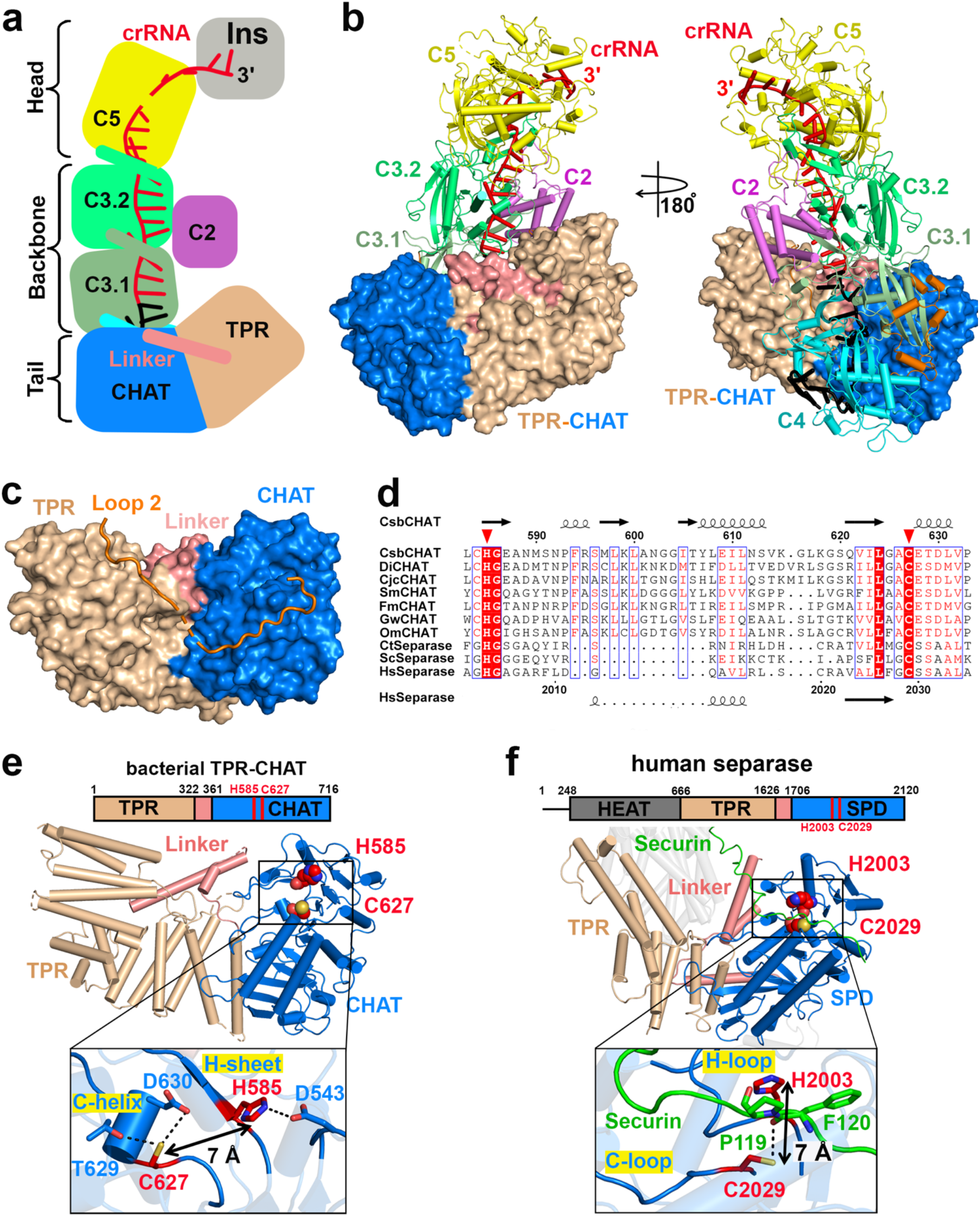
Cryo-EM structure of gRAMP^crRNA^ in complex with TPR-CHAT. **a** and **b**, Schematic 647 (**a**) and ribbon (**b**) representations of cryo-EM structure of gRAMP^crRNA^-TPR-CHAT complex at 648 2.6 Å resolution. The associate TPR-CHAT subunit is shown in surface view. **c**, TPR-CHAT binds to Loop 2 in gRAMP. **d**, Multiple sequence alignment of representative CHAT and separase orthologues. The conserved catalytic residues H585 and C627 are indicated by red triangles. CsbCHAT: KHE91663.1; DiCHAT: WP_124327588.1; CjcCHAT: KAA0249747.1; SmCHAT: OBJA01001127.1; FmCHAT: SESD01000293.1; GwCHAT: MGTA01000040.1; OmCHAT: PDWI01005922.1; Ct: *Chaetomium thermophilum*; Sc: *Saccharomyces cerevisiae*; Hs: *Homo sapiens*. **e** and **f**, Structures of bacterial TPR-CHAT (**e**) and human separase (PDB 7NJ1) (**f**) with an expanded view of the catalytic pocket. The catalytic residues are shown as red spheres. TPR, Tetratricopeptide Repeat Domain; CHAT, Caspase HetF Associated with TPRs; SPD, Separase Protease Domain.

### Structure of bacterial TPR-CHAT resembles eukaryotic separase

A DALI structural comparison revealed TPR-CHAT structurally resembles eukaryotic separase, which is in line well with previous bioinformatic analysis that the bacterial CHAT domain is most closely related to the eukaryotic separase, a caspase-like cysteine protease essential for cohesion dissolution during chromosome segregation^33^. In addition, the bacterial TPR-CHAT contains the highly conserved histidine-cysteine catalytic dyad typical of caspase-like proteases across bacteria and humans (Fig. 3d), implicating that the CHAT protease domain might be a bacterial precursor of eukaryotic separase. In eukaryotes, the separase activity is tightly regulated, otherwise it would lead to missegregation and aneuploidy, thereby resulting in birth defects and cancer^34–36^. The separase activity is typically inhibited by forming a stable complex with securin, while it is activated by degradation of the inhibitory securin^36^.

The structural similarity between bacterial TPR-CHAT and human separase including the highly conserved catalytic dyad H585 and C627 (corresponding to H2003 and C2029 in human separase) (Fig. 3e, f), suggestive of a potential protease activity of bacterial TPR-CHAT. A recent publication has indicated the bacterial TPR-CHAT might be able to cleave and activate bacterial cell death effector gasdermin, executing cell death during antiphage defense^23^, suggestive that the bacterial TPR-CHAT protease might be involved in preventing bacteriophage propagation by host suicide, thereby requiring that its activity also be strictly regulated in bacteria. Notably, lacking an inhibitory peptide, we observed the catalytic C627 and H585 in TPR-CHAT are located in a rigid α helix and a β sheet, named as C-helix and H-sheet, and buried by its own residues (D543, T629 and D630) via electrostatic contacts, leading to inaccessibility of the catalytic residues to incoming substrates. In addition, the distance between H585 and C627 in bacterial TPR-CHAT is ~7 Å (Fig. 3e, insert), a distance similar to that between the H2003 and C2029 in the inactive human separase (Fig. 3f, insert), the activity of which is inhibited by forming a complex with the inhibitory securin, suggesting the TPR-CHAT adopts an inactive state. Taken together, we speculate that TPR-CHAT protease should adopt an autoinhibitory inactivated conformation by forming a stable complex with gRAMP^crRNA^, reminiscent of the autoinhibited ssDNase activity of Cas10 subunit in type III-A Csm^crRNA^ complex^24,25,32^.

### Structural rearrangement of TPR-CHAT induced by the invasive non-self RNA target binding

To figure out if the TPR-CHAT protease activity could be activated during phage infection, we first investigated the conformational changes of TPR-CHAT upon binding of invasive non-self RNA target to the gRAMP^crRNA^-TPR-CHAT complex. We then determined a 2.9 Å cryo-EM structure of an invasive non-self RNA target bound gRAMP^crRNA^-TPR-CHAT quaternary complex (Fig. 4a-c, and Extended Data Fig. 8a-c). The overall structure adopts a similar architecture to the gRAMP^crRNA^-TPR-CHAT ternary complex with an r.m.s.d of 1.78 Å, containing a 38-nt crRNA and 23-nt traceable segment of 56-nt non-self RNA target (Fig. 4a), but with a significant conformational rearrangement of Linker and CHAT protease domains of TPR-CHAT by as much as ~26 Å (Fig. 4d).

**Fig. 4.**
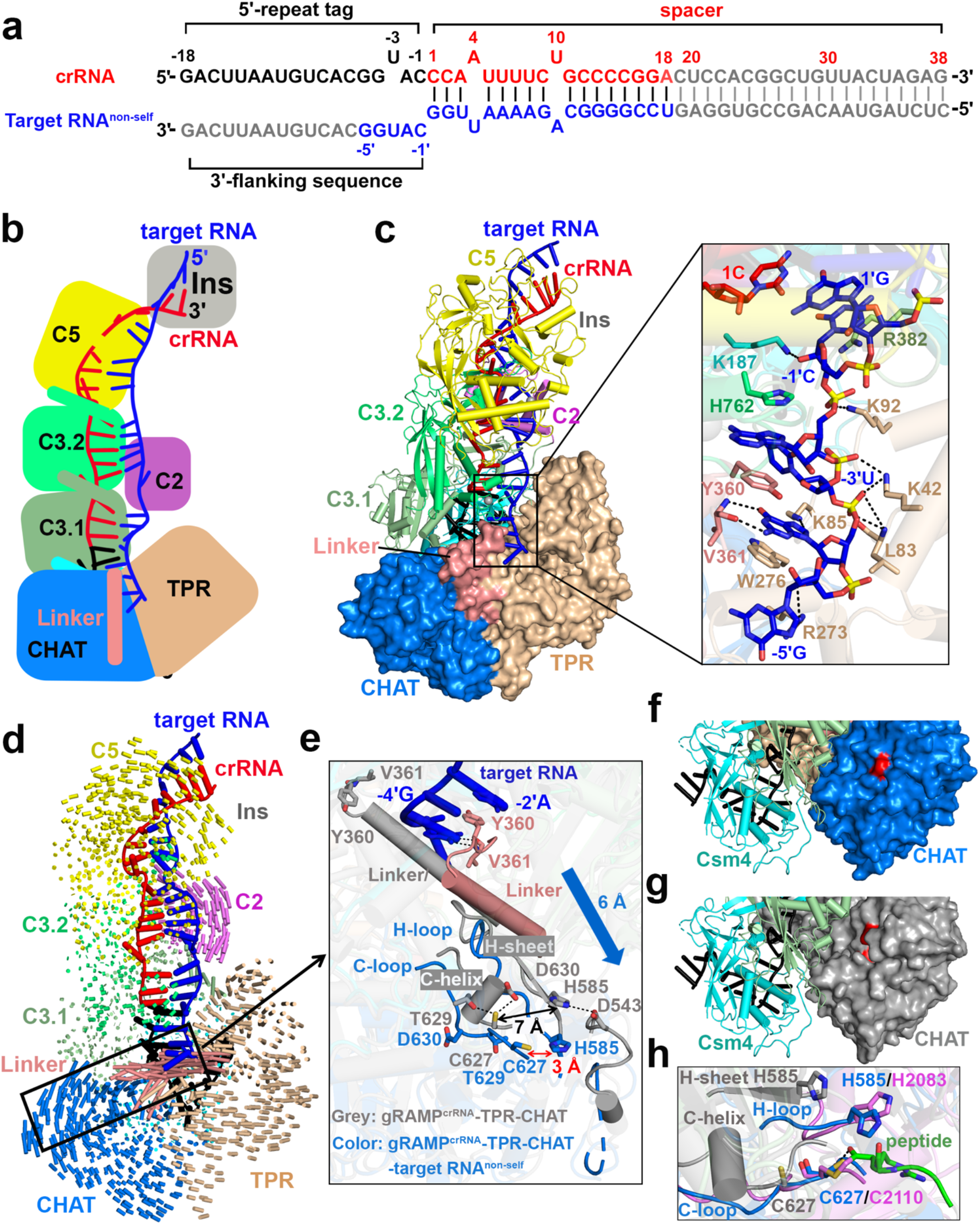
Conformational changes of gRAMP^crRNA^ -TPR-CHAT upon binding of invasive non-self RNA target. **a,** Schematic drawing for the sequences of crRNA and non-self RNA target. Segments that can be traced are in color, while disordered segments are in grey. **b** and **c**, Schematic (**b**) and ribbon (**c**) representations of cryo-EM structure of gRAMP^crRNA^-TPR-CHAT complex bound to non-self RNA target. TPR-CHAT protein is shown in a surface view. The expanded view shows the detailed interactions between 3’-flanking sequence and protein residues in gRAMP and TPR-CHAT. **d**, Structural comparison between gRAMP^crRNA^-TPR-CHAT complex before and after binding to non-self RNA target. Vector length correlates with the domain movement scale. **e**, Detailed conformational changes induced by non-self target RNA binding. The gRAMP^crRNA^-TPR-CHAT complex is shown in grey, and gRAMP^crRNA^-TPR-CHAT-target RNA^non-self^ complex is shown in color. The blue arrow indicates the movement scale of the secondary structures in the CHAT protease catalytic pocket. **f** and **g**, Structures of CHAT protease domain in gRAMP^crRNA^ -TPR-CHAT before (**g**) and after (**f**) non-self RNA target binding. The CHAT domain is shown in a surface view. The red surface indicates the position of catalytic residues H585 and C627. **h**, Structural comparison of the catalytic pockets among gRAMP^crRNA^-TPR-CHAT (in grey), gRAMP^crRNA^-TPR-CHAT-target RNA^non-self^ (in marine) and the active separase protease domain from the thermophilic fungus *Chaetomium thermophilum* (in violet, PDB 5FC3). The substrate peptide of the separase is shown in green.

Notably, the 3’-flanking sequence of the invasive non-self RNA target at position −5’ to −1’ is non-complementary to the 5’-repeat tag of crRNA (Fig. 4a), swinging away from crRNA and directly interacting with TPR-CHAT, where it is lying in a cleft between the Linker and TPR domains (Fig. 4b, c). Nucleotide-1’ C is stabilized by K187^Csm4^ and R382^Csm3.1^, while nucleotides at positions −2’ to −5’ directly interreact with TPR-CHAT mainly through sequence-nonspecific contacts, which induce the large conformational changes of Linker and CHAT protease catalytic pocket (Fig. 4d). Especially, nucleotide −2’ A and −4’ G directly interact with Y360 and V361 of the Linker domain, inducing dramatic conformational changes of Linker domain followed by the rearrangement of H-helix and C-sheet into the extended loop structures in the CHAT protease catalytic pocket (Fig. 4e). The conformational changes in the catalytic pocket result in outward-movement and exposure of the catalytic residues H585 and C627 to be accessible to the potential substrate peptide (in marine, Fig. 4e, f), compared to structure before the non-self RNA target binding (in grey, Fig. 4e, g). Consistently, both the exposure of catalytic cysteine to solvent and the reorganization of C-loop have been shown to indicates the activation of the caspase-like protease^37,38^, suggestive of the activation of TPR-CHAT protease upon invasive non-self RNA target binding. Most importantly, the distance between the catalytic dyad H585 and C627 moves from ~7 Å in inactive gRAMP^crRNA^-TPR-CHAT complex (in grey) to ~3 Å (in marine) upon the non-self RNA target binding, a distance where both these catalytic residues are positioned for attack on the peptide bond of the substrate (Fig. 4e). Furthermore, upon target binding of the invasive non-self RNA, the conformation of C-loop and the geometry of catalytic dyad H585 and C267 switches from an inactive state (in grey) to a state (in marine) similar to the active *Chaetomium thermophilum* separase (in violet)^34^ (Fig. 4h), suggesting that invasive non-self RNA binding allosterically activates the TPR-CHAT protease activity in gRAMP^crRNA^-TPR-CHAT complex.

### The 3’-anti-tag of self RNA binds to a channel distinct from the 3’-flanking sequence of non-self RNA

If binding of the invasive non-self RNA activates the TPR-CHAT protease activity, which might be involved in host suicide, binding of self RNA should not activate the TPR-CHAT protease activity to prevent self-targeting. To test this hypothesis, we have determined a 2.7 Å cryo-EM structure of gRAMP^crRNA^-TPR-CHAT bound to a self RNA target with an anti-tag sequence complementary to 5’-repeat tag (Fig. 5a-c and Extended Data Fig. 8d-f). Superposition of structures between bound self and non-self RNA targets reveals pronounced conformational differences in the positions of Linker and CHAT protease domain (Fig. 5d), indicating gRAMP^crRNA^-TPR-CHAT complex distinguishes self from no-self RNA targets, which is manifested in the distinct conformation of TPR-CHAT.

**Fig. 5.**
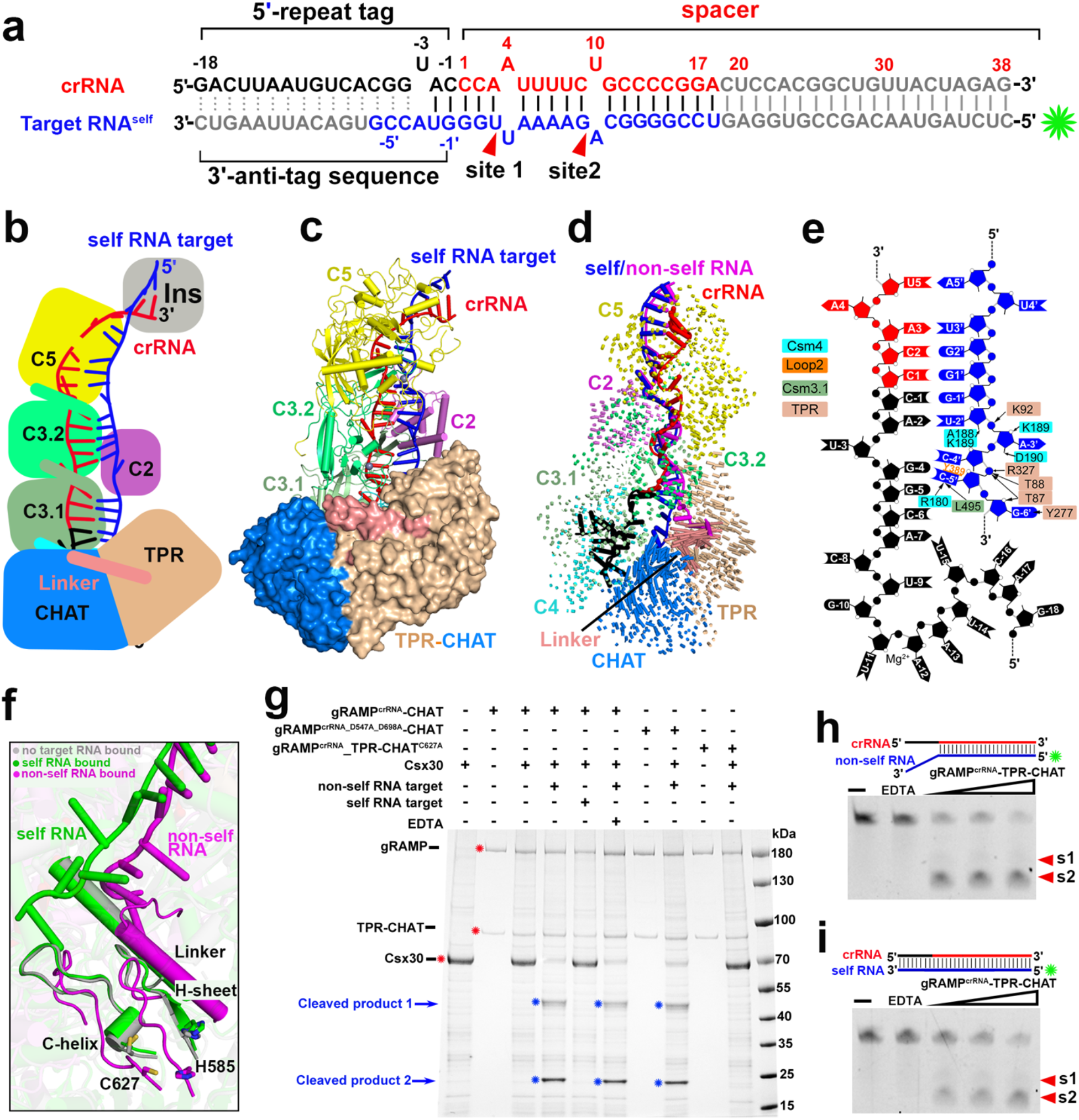
Cryo-EM structure of gRAMP^crRNA^-TPR-CHAT bound to self RNA target. **a**, Schematic representation for the sequences of crRNA and self RNA target. Segments that can be traced are in color, while disordered segments are in grey. **b** and **c**, Schematic (**b**) and cartoon (**c**) representations of cryo-EM structure of self RNA target bound gRAMP^crRNA^-TPR-CHAT quaternary complex. TPR-CHAT protein is shown in a surface view. **d**, Structural comparison between gRAMP^crRNA^-TPR-CHAT complex bound to self and non-self RNA targets. Vector length correlates with the domain movement scale. **e**, Schematic interactions between 3’-flanking sequence and protein residues in gRAMP and TPR-CHAT. Hydrogen bonds are indicated by black arrows, and the stacking interactions are shown by orange lines. The protein residues responsible for the interactions with the 3’-anti-tag of the RNA target are colored according to the domain. **f**, Conformational changes of the Linker and CHAT protease catalytic pocket in TPR-CHAT induced by self (in green) and non-self (in violet) RNA target binding. The TPR-CHAT subunit in gRAMP^crRNA^-TPR-CHAT before target RNA binding is colored in grey. g, Cleavage of Csx30 by the non-self RNA target activated protease in gRAMP^crRNA^-TPR-CHAT complex. The red asterisk indicates the position of input proteins labeled in left panel. The blue asterisk and arrow indicate the cleaved products. **h** and **i**, Cleavage of non-self (**h**) and self (**i**) RNA targets by gRAMP^crRNA^-TPR-CHAT complex. The green asterisk indicates the fluorescence label at 5’ end of the RNA target.

Notably, distinct from 3’-flanking sequence of non-self RNA that lies in the cleft between Linker and TPR domain in TPR-CHAT (Fig. 4c), the 3’-anti-tag sequence of self RNA binds in a channel between TPR-CHAT and gRAMP (Fig. 5c). Nucleotides −1’G and −2’U of self RNA base pair with the nucleotides −1C and −2A within the 5’-repeat tag of crRNA, and these base-pairing contacts drag the other anti-tag sequence at position −3’ to −6’ to bind in a cleft between Csm4 and TPR domain via both sequence-specific and -nonspecific contacts (Fig. 5e). This binding mode of 3’-anti-tag sequence of self RNA induces minimal conformational changes of the gRAMP^crRNA^-TPR-CHAT complex, including the conformation of C-helix and H-sheet and the geometry of catalytic dyad H585 and C267 in the CHAT protease domain (Fig. 5f and Extended Data Fig. 9a), which contrasts with those induced by 3’-flanking sequence of non-self RNA (Fig. 4d and 5g), suggesting that TPR-CHAT protease still adopts an autoinhibitory conformation upon own self RNA target binding, thereby preventing self-targeting to avoid autoimmunity.

To test if only foreign non-self RNA target could activate the TPR-CHAT protease activity in vitro, we first have to find the substrates of the TPR-CHAT protease. Previous reports have shown that the substrates of CHAT proteases are commonly encoded by genes next to the protease-domain containing genes^23^. In type III-E CRISPR loci (Fig. 1a), except the gene is known to encode the sigma factor RpoE, the other two genes encoding proteins Csx30 and Csx31 remain mysterious. We then wondered whether Csx30 and Csx31 could be the substrates of TPR-CHAT protease. As shown in Fig. 5g and Extended Data Fig. 9b and c, only Csx30 could be cleaved into two separate products by gRAMP^crRNA^-TPR-CHAT complexes upon non-self RNA target binding but not self RNA target binding, which in line well with our structural analysis. Besides, mutation of the catalytic residue C267 in TPR-CHAT protease into alanine abolished the protease cleavage activity (Fig. 5g), further demonstrating the catalytic pocket in TPR-CHAT protease. As expected, neither mutation of the RNase catalytic residues nor addition of EDTA, which suppress RNase activities of Csm3 domains (Fig. 2c, d), affected TPR-CHAT protease activities in gRAMP^crRNA^-TPR-CHAT complexes (Fig. 5g). Noteworthy, both non-self and self RNA targets could be recognized and cleaved by Csm3 subunits in gRAMP^crRNA^-TPR-CHAT complex (Fig. 5h, i), suggesting that the RNase activities provides a temporal control of CHAT protease activities and thereby preventing potential damage of the continuous enzymatic activities to host cells. Thus, the base-paring potential between crRNA and target RNA at −1 and −2 position, rather than the RNase activities, is responsible for virus-host discrimination and determines whether the associated TPR-CHAT protease activity could be switched on or stay off, providing an ancient regulation mechanism for the caspase-like protease activity distinct from its eukaryotic homolog separase.

## Discussion

Type III-A CRISPR-Cas immunity against invading bacteriophages and plasmids is achieved by crRNA-bound multi-subunit Csm complex (Csm2 to Csm5, and Cas10 subunits) by binding to the invasive non-self RNA target, triggering the activation of ssDNase and cyclic oligoadenylate (cOA) synthetase activities of the signature protein Cas10^19,20^ (Fig. 6a, b). The produced second messenger cOA allosterically activates the non-specific RNase Csm6, which provides host with further protection against invading phages and plasmids^39,40^. Both enzymic activities of Cas10 are switched off by sequence-specific cleavage of the invasive RNA targets by the RNase Csm3 subunits, providing a temporal control of Cas10 activities and thereby preventing potential damage to host cells. The base-pairing potential between elements of the 5’-repeat tag at position −2 to −5 and 3’-sequence of target RNA is responsible for discriminating self from non-self RNA targets^2,18^.

**Fig. 6.**
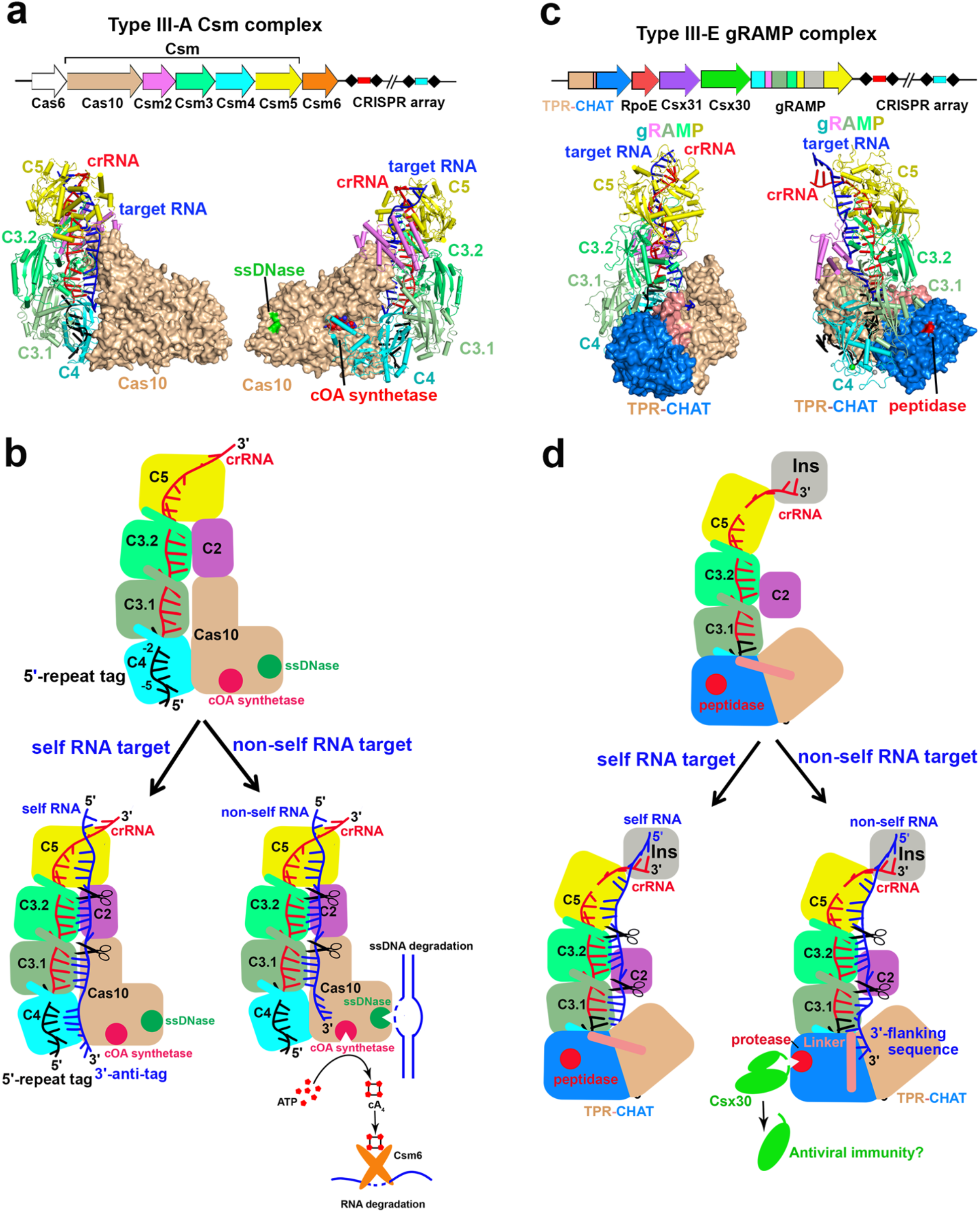
Structural and mechanistic comparisons between type III-A and type III-E effector 702 complexes. **a** and **c**, Overall architecture of the effector complex and CRISPR-locus in type III-A 703 (**a**) and type III-E systems (**c**). The ssDNase and cOA synthesis pocket in type III-A effector complex are colored in green and red, respectively. The protease pocket in type III-E effector complex is colored in red. **b**, Model of type III-A interference during anti-phage or -plasmid defense. The type III-A muti-subunit effector complex uses crRNA to recognize the invasive non-self RNA targets with 3’-flanking sequence noncomplementary to 5’-repeat tag, which triggers the non-specific cleavage of ssDNA and synthesis of the second messenger cyclic oligoadenylate cOA by the signature Cas10 protein. The produced cOA binds to Csm6 and activates its single-strand RNase activity, which are crucial for type III-A immunity in vivo. Both the enzymic activities of Cas10 are switched off upon the subsequent cleavage of non-self RNA targets at 6-nt intervals by Csm3 subunits. In contrast, the self RNA containing an 3’-anti-tag sequence complementary to 5’-repeat tag could not activates Cas10 enzymatic activities, but still could be cleaved by Csm3 subunits. **d**, Model of type III-E interference during anti-phages or -plasmids infection. The type III-E single -protein effector forms a stable complex with a caspase-like protease TPR-CHAT, which adopts an autoinhibitory state. Binding of non-self RNA targets with 3’-flanking sequence noncomplementary to 5’-repeat tag introduces significant conformational changes of the TPR-CHAT linker domain, inducing rearrangement of the protease catalytic domain. These conformational changes induced by 3’-flanking sequence of non-self target RNA activates the protease activity of the CHAT domain, leading to cleavage of target Csx30 proteins, which might be involved in the antiphage defense. Cleavage of target RNA into 6-nt intervals by Csm3 domains would shut down the protease activity. The 3’-anti-tag sequence takes a different binding route upon base-paring with 5’-repeat tag of crRNA, which could not induce conformational changes of the Linker and CHAT protease domains, thus leaving the protease to remain in an autoinhibitory inactive state. The bound self RNA targets could still be cleaved by Csm3 domains.

A recently reported type III-E CRISRP-Cas system contains a single-protein effector, demonstrated specific recognition and cleavage of target RNA without collateral activity and cell toxicity^12,13^, thereby representing a new promising RNA manipulation tool. Most intriguingly, lacking the signature Cas10 protein crucial for canonical type III CRISPR immunity, type III-E CRISPR loci contains a gene encoding a caspase-like protease TPR-CHAT, which forms a stable complex with type III-E effector complex gRAMP, and sits in a similar position as Cas10 does in type III-A effector complex (Fig. 6a, c). The caspase-like protease TPR-CHAT has been implicated in regulating cell death during antiphage defense^23^, raising the intriguing possibility that the type III-E CRISPR-caspase systems might defend viral infection by host suicide. Using structural biology and biochemistry, we have provided the first glimpse of the structural basis for the functional relationship between caspase-like protease and CRISPR-Cas systems, leading to a proposed structural model for the target RNA-activated TPR-CHAT protease activity to reach type III-E antiviral immunity, thereby suggesting a neofunctionalization of type III-E CRISPR-Caspase systems (Fig. 6d).

TPR-CHAT protease stays in an inactive autoinhibitory state by forming a stable complex with gRAMP^crRNA^. Upon binding of invasive non-self RNA targets, its 3’-flanking sequence induces the conformational changes of the TPR-CHAT Linker domain by as much as ~20 Å followed by the rearrangement of the CHAT protease catalytic pocket, leading to the activation of the CHAT protease that cleaves the protein substrate Csx30. The autoinhibitory rigid C-helix and H-sheet in the catalytic pocket are transformed into flexible C-loop and H-loop, while the geometry of catalytic dyad H585 and C267 become accessible for attacking the peptide bond of the substrate peptide (Fig. 4e, h). The subsequent cleavage of the RNA targets by Csm3 domains would release the RNA targets, thereby switching off the TPR-CHAT protease activity (Fig. 5g, h). By contrast, for binding of self RNA target, the 3’-anti-tag sequence takes a route distinct from 3’-flanking sequence of non-self RNA target, while inducing minimal conformational changes of the TPR-CHAT protease (Fig. 5f, and Extended Data Fig. 9a), which still remains in an autoinhibitory state. Thus, the base-paring potential between elements of the 5’-repeat tag of crRNA and 3’-sequence of RNA target at position −1 and −2 plays a key role in discriminating self from non-self (Fig. 4a, 5a). The RNase activity of Csm3 domains possibly just contributes to temporal control of TPR-CHAT protease activity, thereby preventing potential damage to host cells.

An interesting question that remains to be addressed relates to whether and how the cleaved Csx30 could induced the bacterial death, given that it has been implicated that TPR-CHAT protease has the ability to cleave and activate bacterial cell death effector gasdermin, executing cell death during antiphage defense^23^. Our studies have established structural homology between bacterial TPR-CHAT protease and eukaryotic separase, which is known to cleave after the conserved EXXR (X represents any residue) peptide motif^34^. However, the critical separase residue D2151 responsible for the conserved R215 recognition at the P1 position is replaced by K670 in bacterial TPR-CHAT, and the residues around E212 at P4 position is also different between eukaryotic separase and bacterial TPR-CHAT (Extended Data Fig. 9d), suggesting a distinct substrate specificity of TPR-CHAT that is distinct from that for eukaryotic separase^34^. It will be of interest to investigate the specific cleavage sites of Csx30 by TPR-CHAT protease and its potential involvement in the CRISPR-mediated defense pathways in future studies. Nevertheless, our findings have elucidated RNA-targeting mechanisms by the gRAMP^crRNA^ complex, providing a structural basis for the development of the new promising gRAMP^crRNA^-based RNA manipulation tools. In addition, we have provided the first insights into the structural basis for the functional relationship between type III-E CRISPR-gRAMP effector complex and caspase-like TPR-CHAT protease. Furthermore, we provide a mechanism for self versus non-self RNA target discrimination governed by conformational changes-induced activation of TPR-CHAT protease in type III-E CRISPR-Caspase antiviral immunity.

## Methods

### Protein expression and purification

The full-length *Candidatus* “Scalindua brodae” genes encoding gRAMP, TPR-CHAT, RpoE, Csx30, and Csx31 were synthesized by Sangon Biotech (Shanghai). The *csx30* gene was cloned into the pCDFDuet-1 (Novagen, streptomycin resistance) vector with an N-terminal hexahistidine tag and a C-terminal Strep II tag. The *csx31* gene was cloned into a modified pET28a vector with a SUMO tag following a ubiquitin-like protease (ULP1) cleavable site and an MBP tag to increase the soluble protein yield, fused with an N-termial hexahistidine tag. The gRAMP gene and the synthetic CRISPR gene were cloned into the pRSFDuet-1 vector (Novagen, kanamycin resistance), in which an N-terminal hexahistidine tag was fused with gRAMP. The TPR-CHAT gene was subcloned into the pETDuet-1 vector (Novagen, ampicillin resistance) with a C-terminal Strep II tag. Individual vector or two vectors were transformed into *Escherichia coli* BL21-CodonPlus (DE3). Streptomycin-resistant colonies for expression of Csx30, kanamycin-resistant colonies for expression of Csx31, double-resistant colonies for expression of gRAMP^crRNA^-TPR-CHAT complex or kanamycin-resistant colonies for expression of gRAMP^crRNA^ complex were picked and grown to OD_600_ of ~ 0.8 in LB medium at 37°C. The expression was induced by adding isopropyl-b-D-1-thiogalactopyranoside to 0.5 mM and shifted to at 16 °C for 20 hrs. Cells were harvested by centrifugation and resuspended in lysis buffer (20 mM Tris-HCl, 300 mM NaCl, 20 mM imidazole, 1 mM DTT, pH 8.0). The harvested cells were then lysed by the EmulsiFlex-C3 homogenizer (Avestin) and centrifuged at 20,000 rpm for 30 min.

For the purification of gRAMP^crRNA^ complex or Csx31, the supernatant was applied to 5 mL HisTrap Fast flow column (Cytiva). The protein was eluted with lysis buffer supplemented with 500 mM imidazole after washing the column with 10 column volumes of lysis buffer and 2 column volumes of lysis buffer supplemented with 40 mM imidazole.

For the purification of the gRAMP^crRNA^-TPR-CHAT complex or Csx30, the elution from HisTrap Fast flow column was subjected to further purification by loading into 5 mL StrepTrap HP column (Cytiva). Proteins were eluted in buffer A (20 mM Tris-HCl, 100 mM NaCl, 1 mM DTT, pH 8.0) supplemented with 5 mM desthiobiotin and then loaded into 5 mL HiTrap Q Fast Flow column (Cytiva). Proteins were eluted in a linear gradient from 100 mM to 1 M NaCl in 20 column volumes, and then concentrated using 100 kDa molecular mass cut-off concentrators (Amicon) before final purification on a Superdex 200 increase 10/300 GL column (Cytiva) pre-equilibrated in buffer B (20 mM Tris-HCl, 150 mM NaCl, 1 mM DTT, pH 8.0). Pooled fractions were concentrated, flash frozen in liquid nitrogen and stored at −80°C for further use.

All mutants were generated by site-directed mutagenesis, and purified by the same method as above.

### In vitro assembly of gRAMP^crRNA^ or gRAMP^crRNA^-TPR-CHAT in complex with self or non-self RNA targets

To assemble gRAMP^crRNA^-target RNA or gRAMP^crRNA^-TPR-CHAT-target RNA ternary complex, the purified gRAMP^crRNA^ binary or gRAMP^crRNA^-TPR-CHAT ternary complex were mixed with either self RNA (5’ CUCUAGUAACAGCCGUGGAGUCCGGGGCAGAAAAUUGGGUACCGUGACAUUAAG UC 3’) or non-self RNA (5’ CUCUAGUAACAGCCGUGGAGUCCGGGGCAGAAAAUUGGCAUGGCACUGUAAUUC AG 3’) at a molar ration of 1:1.2 and then incubated on ice for 30 min. We have added 5 mM EDTA to chelate divalent cations and thus prevent target cleavage. The mixture was loaded on a Superdex 200 Increase 10/300 GL(Cytiva) column equilibrated with buffer B. Pooled fractions were concentrated, flash frozen in liquid nitrogen and stored at −80°C.

### Cryo-EM sample preparation and data acquisition

3.5 μL of 2 mg/mL purified gRAMP^crRNA^, gRAMP^crRNA^-target RNA^self^, gRAMP^crRNA^-target RNA^non-self^, gRAMP^crRNA^-TPR-CHAT, gRAMP^crRNA^-TPR-CHAT-target RNA^self^ and gRAMP^crRNA^-TPR-CHAT-target RNA^non-self^ complexes were applied onto glow-discharged UltrAuFoil 300 mesh R1.2/1.3 grids (Quantifoil), respectively. Grids were blotted for 2 s at 100% humidity, 4°C and flash frozen into liquid ethane using Vitrobot Mark IV (FEI). Images were collected on the FEI Titan Krios electron microscope at acceleration voltage of 300 kV with a Gatan K3 Summit detector with a physical pixel size of 1.1 Å at a defocus range from −1.5 μm to −2.5 μm. Each movie comprises 32 subframes with a total dose of 50 e^−^/Å^2^.

### Cryo-EM data processing

Image processing were performed by RELION 3.1 ^41^ and cryoSPARC v3.1 ^42^. The motion correction was performed with MotionCor2 ^43^. Contrast transfer function (CTF) parameters were estimated by Ctffind4 ^44^. Auto-picked particles using Laplacian-of-Gaussian were extracted and subjected to two rounds of 2D classification and two rounds of 3D classification in cryoSPARC v3.1, using the initial model generated in cryoSPARC v3.1 as a reference. Particles corresponding to the best class with the highest-resolution features were selected and subjected to non-uniform refinement in cryoSPARC v3.1. The final particles were subjected to the CTF refinement and then subjected to non-uniform refinement, generating the final reconstitutions. To further improve the density of the Csm5 insertion domain in the gRAMP^crRNA^-target RNA^self^ complex, a soft mask for the insertion domain was generated and applied for the subsequent local refinement in cryoSPARC v3.1. The final map of gRAMP^crRNA^-target RNA^self^ complex was obtained by combining the maps from non-uniform refinement and local refinement in Chimera ^45^. All resolutions were estimated by applying a soft mask around the protein density and the Fourier shell correlation (FSC) = 0.143 criterion in cryoSPARC v3.1. Local resolution estimates were calculated from two half data maps in cryoSPARC v3.1. Further details related to data processing and refinement are summarized in Extended Data Table 1.

### Atomic model building and refinement

Atomic models were built de novo, then interactively refined in COOT ^46^. All models were refined against the cryo-EM maps using phenix.real_space_refine ^47^ by applying geometric and secondary structure restraints. All figures were prepared in PyMol (http://www.pymol.org) and Chimera ^45^. The statistics for data collection and model refinement are shown in Extended Data Table 1.

### RNA cleavage assay

In vitro RNA cleavage assays were performed by incubating gRAMP^crRNA^ complex or its mutants in different concentrations as indicated with 50 nM 5’ 6-FAM labelled RNA in 10 μL reaction buffer composed of 20 mM HEPES, 75 mM NaCl, 5 mM MgCl_2_, 100 μM CoCl_2_, pH 7.5 at 37°C for 60 min. The reactions were quenched by the addition of protease K and 5 × stop buffer (0.5 mg/mL bovine serum albumin, 5% SDS, 50 mM EDTA) followed by incubation for 10 mins at 95°C. The samples were separated on 15% 8 M urea-PAGE and the FAM fluorescence signal was quantified with Canon 6100 Multi imaging system (Tanon Science & Technology Co., Ltd., Shanghai, China). The cleavage assays were repeated at least three times, and representative results are shown.

### Protease cleavage assay

The cleavage reactions were assembled in 10 μL reaction volume with buffer containing 20 mM Tris-HCl, 100 mM NaCl, pH 8.0. 1μM gRAMP^crRNA^-TPR-CHAT complex or its mutants (gRAMP_D547A _D698A^crRNA^-TPR-CHAT, gRAMP^crRNA^-TPR-CHAT_C627A) or TPR-CHAT protease were added to reactions with 5μM proposed substrates (Csx30 or Csx31), followed by the addition of self RNA or non-self RNA target. Reactions were incubated at 37°C for 60 min, then separated on 4-20% FuturePAGE™ protein gel (ACE). Gels were stained with Coomassie G-250, and imaged using ChampGel7000 (Sage).

## Data availability

The cryo-EM density maps have been deposited in the Electron Microscopy Data Bank (EMDB) under accession number EMD-33676 (gRAMP^crRNA^); EMD-33678 (gRAMP^crRNA^-target RNA^self^); EMD-33677 (gRAMP^crRNA^-target RNA^non-self^); EMD-33680 (gRAMP^crRNA^-TPR-CHAT); EMD-33681 (gRAMP^crRNA^-TPR-CHAT-target RNA^self^) and EMD-33679 (gRAMP^crRNA^-TPR-CHAT-target RNA^non-self^). The atomic coordinates have been deposited in the Protein Data Bank (PDB) with accession number 7Y80 (gRAMP^crRNA^); 7Y82 (gRAMP^crRNA^-target RNA^self^); 7Y81 (gRAMP^crRNA^-target RNA^non-self^); 7Y84 (gRAMP^crRNA^-TPR-CHAT); 7Y85 (gRAMP^crRNA^-TPR-CHAT-target RNA^self^) and 7Y83 (gRAMP^crRNA^-TPR-CHAT-target RNA^non-self^).

## Acknowledgments

We thank the staff at Southern University of Science and Technology (SUSTech) Cryo-EM Center for assistance in data collection on the Titan KRIOS cryo-electron microscope. This research was supported by the Shenzhen Government ‘Peacock Plan’ (Y01416126 to N.J.) and SUSTech start-up funding (Y01416226 to N.J.), the Guangdong Provincial Science and Technology Innovation Council Grant (2017B030301018 to N.J.), the Shenzhen Science and Technology Program (KQTD20190929173906742 to H.H), Key Laboratory of Molecular Design for Plant Cell Factory of Guangdong Higher Education Institutes (2019KSYS006 to H.H), and the Shenzhen Government ‘Peacock Plan’ (Y01226136 to H.H).

## Author contributions

N.C. undertook biochemical and structural studies, from sample preparation and purification, biochemical assays, and cryo-EM data collection. J.T.Z performed data processing, and structure refinement and analysis. In addition, C.W. shared his cryo-EM expertise with J.T.Z. and N.J. to facilitate the research. Z.L. and X.Y.L. contributed to recombinant plasmid construction and protein purification. N.J. and H.H directed the research. N.J. wrote the manuscript with input from other authors.

---

**Extended Data Fig. 1.**
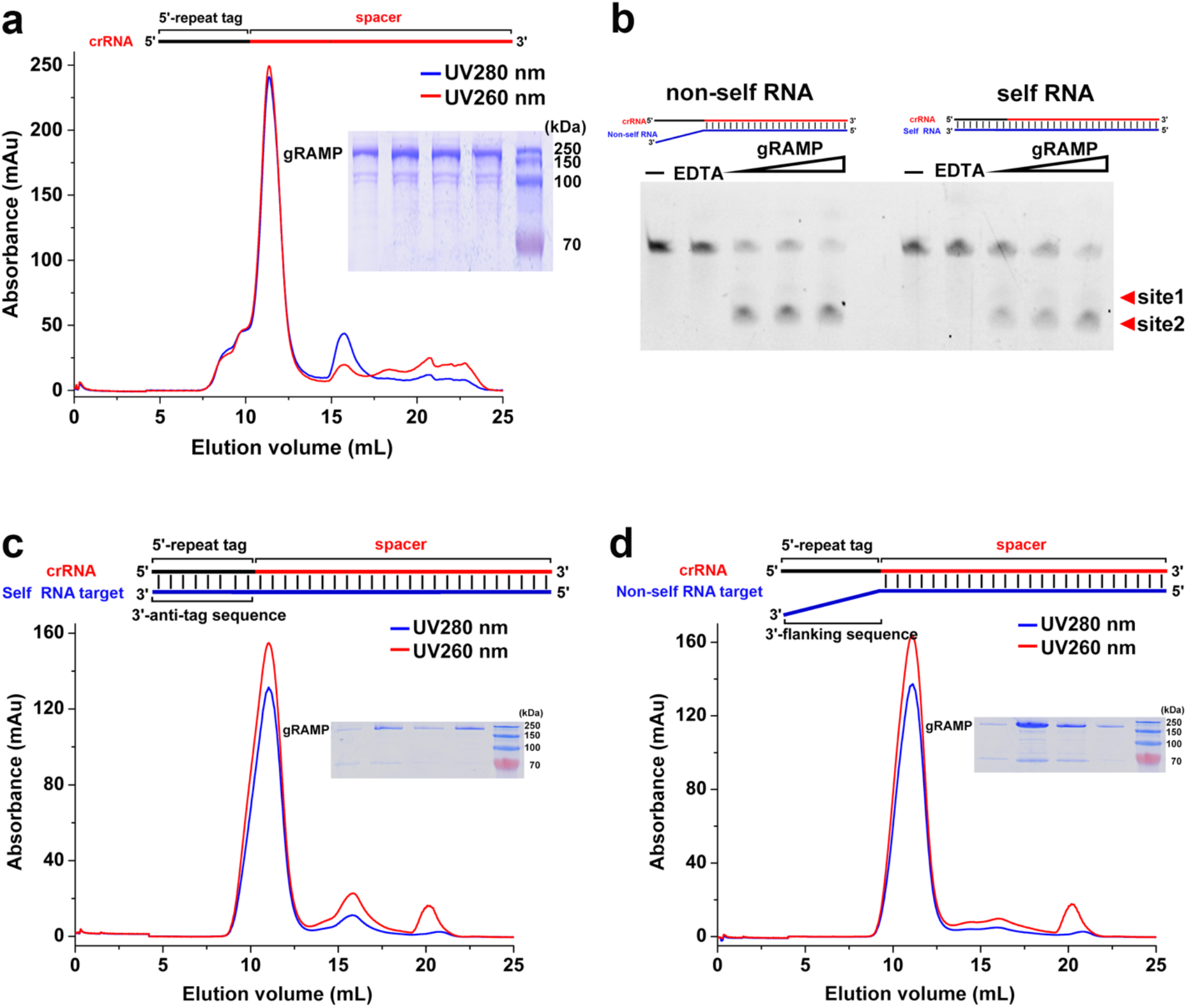
Biochemical Reconstruction of *Candidatus* “Scalindua brodae” gRAMP^crRNA^ binary, gRAMP^crRNA^-target RNA^self^ and gRAMP^crRNA^-target RNA^non-self^ ternary complexes. **a**, Size-exclusion chromatography and SDS-PAGE profile for purification of gRAMP^crRNA^ binary complex. **b**, Cleavage of target RNA by gRAMP^crRNA^ complex. **c** and **d**, Size-exclusion chromatography and SDS-PAGE profile for *in vitro* assembly of gRAMP^crRNA^ complex with either self RNA (**c**) or non-self (**d**) RNA targets. Red and blue curves correspond to 260 and 280 nm UV absorptions, respectively.

**Extended Data Fig. 2.**
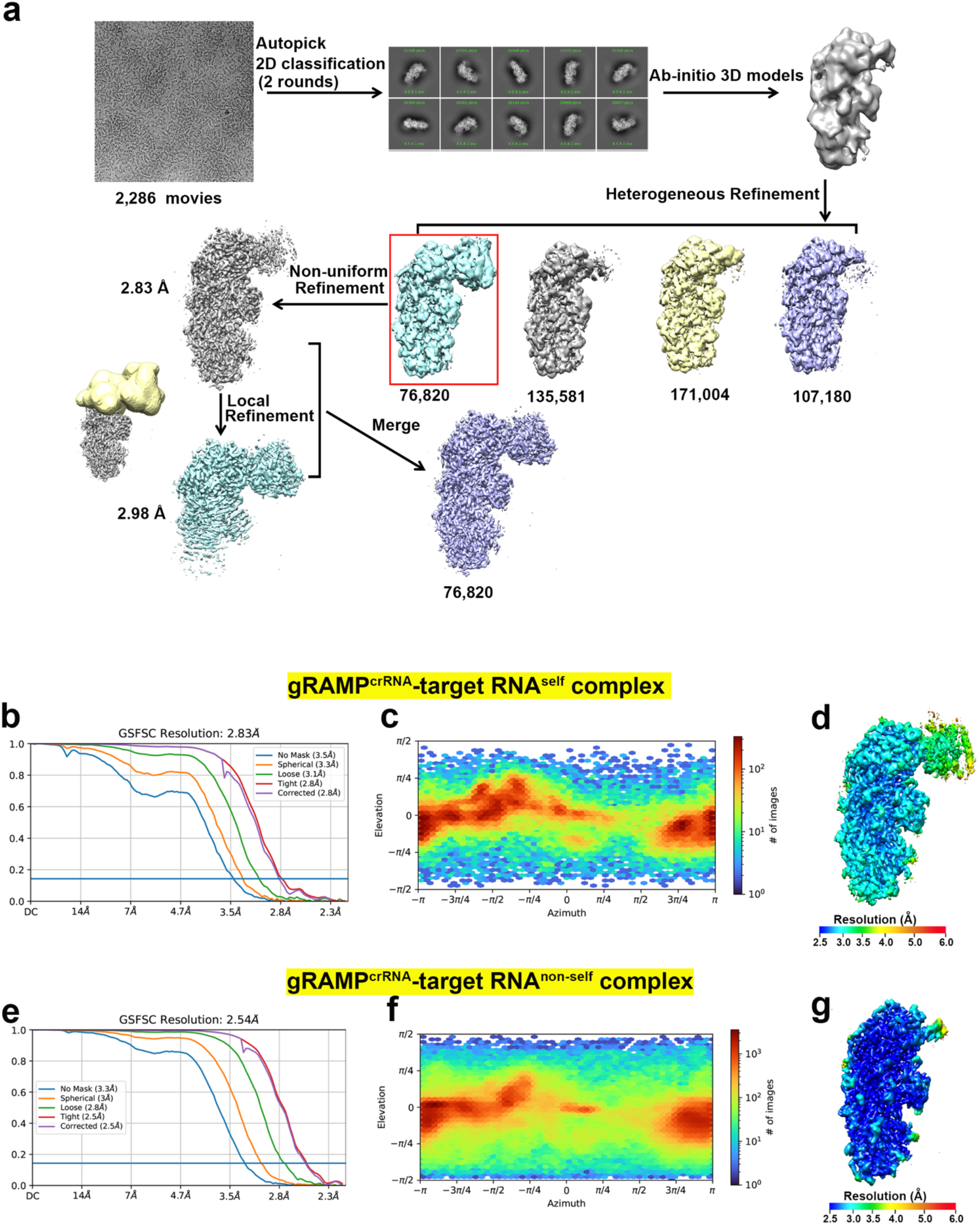
Cryo-EM Reconstruction of gRAMP^crRNA^-target RNA^self^ and 744 gRAMP^crRNA^-target RNA^non-self^ ternary complexes. **a**, Flow chart of image processing for 745 gRAMP^crRNA^-target RNA^self^ ternary complex. **b** and **e**, Fourier Shell Correlation (FSC) curve of gRAMP^crRNA^-target RNA^self^ (**b**) and gRAMP^crRNA^-target RNA^non-self^ (**e**) ternary complex reconstruction. **c** and **f**, Direction distribution plot of gRAMP^crRNA^-target RNA^self^ (**c**) and gRAMP^crRNA^-target RNA^non-self^ (**f**) ternary complex reconstruction. **d** and **g**, Final 3D reconstructed map of gRAMP^crRNA^-target RNA^self^ (**d**) and gRAMP^crRNA^-target RNA^non-self^ (**g**) ternary complex colored according to local resolution.

**Extended Data Fig. 3.**
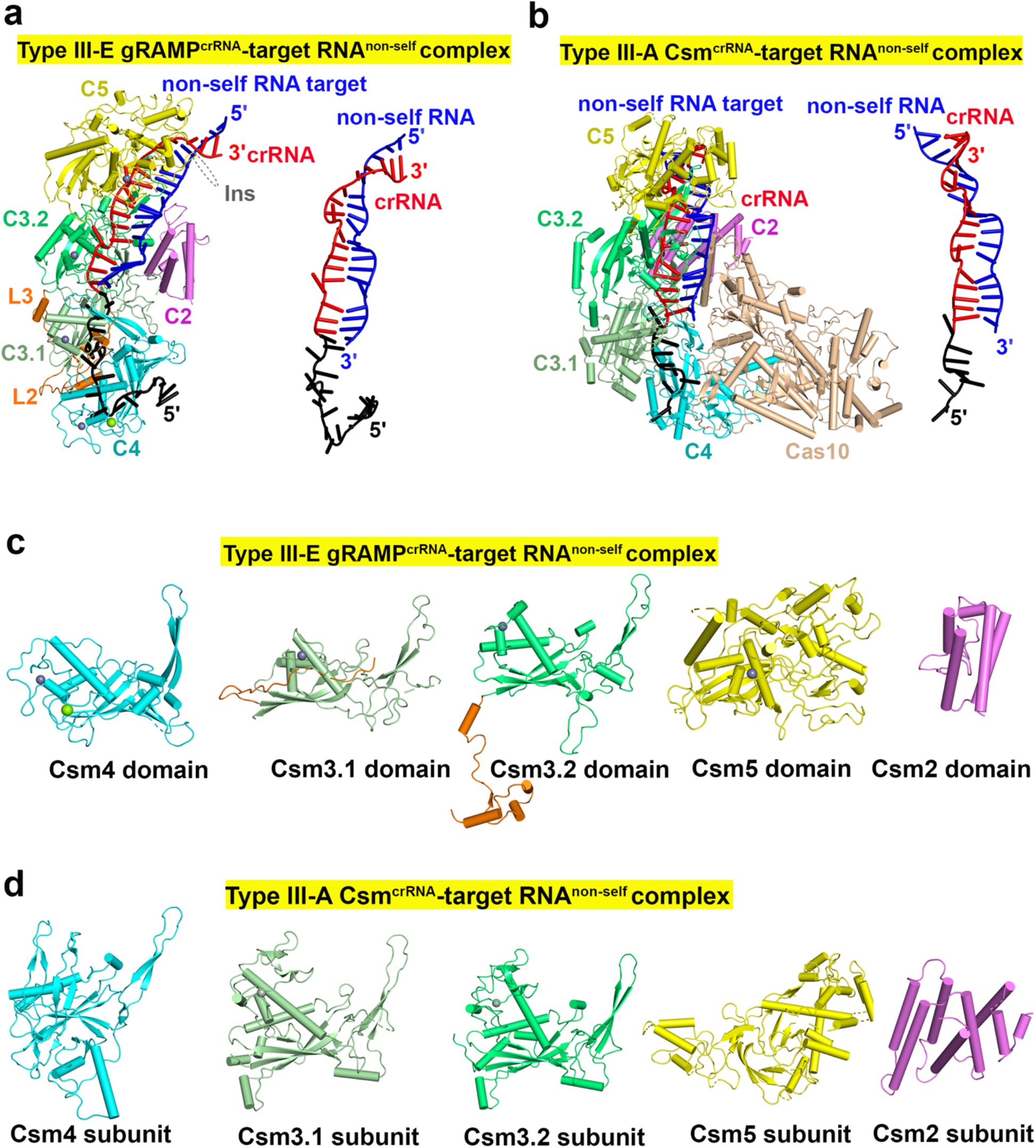
Structural comparison between type III-E gRAMP^crRNA^-target RNA^non-self^ and type III-A Csm^crRNA^-target RNA^non-self^ complexes. **a** and **b**, Cryo-EM structure of type III-E gRAMP^crRNA^-target RNA^non-self^ complex (**a**), Csm^crRNA^-target RNA^non-self^ complex (**b**, PDB 6MUR) and its crRNA-target RNA^non-self^ duplex. **c** and **d**, Individual subunits of the type III-E gRAMP (**c**) and type III-A Csm (**d**) complexes.

**Extended Data Fig. 4.**
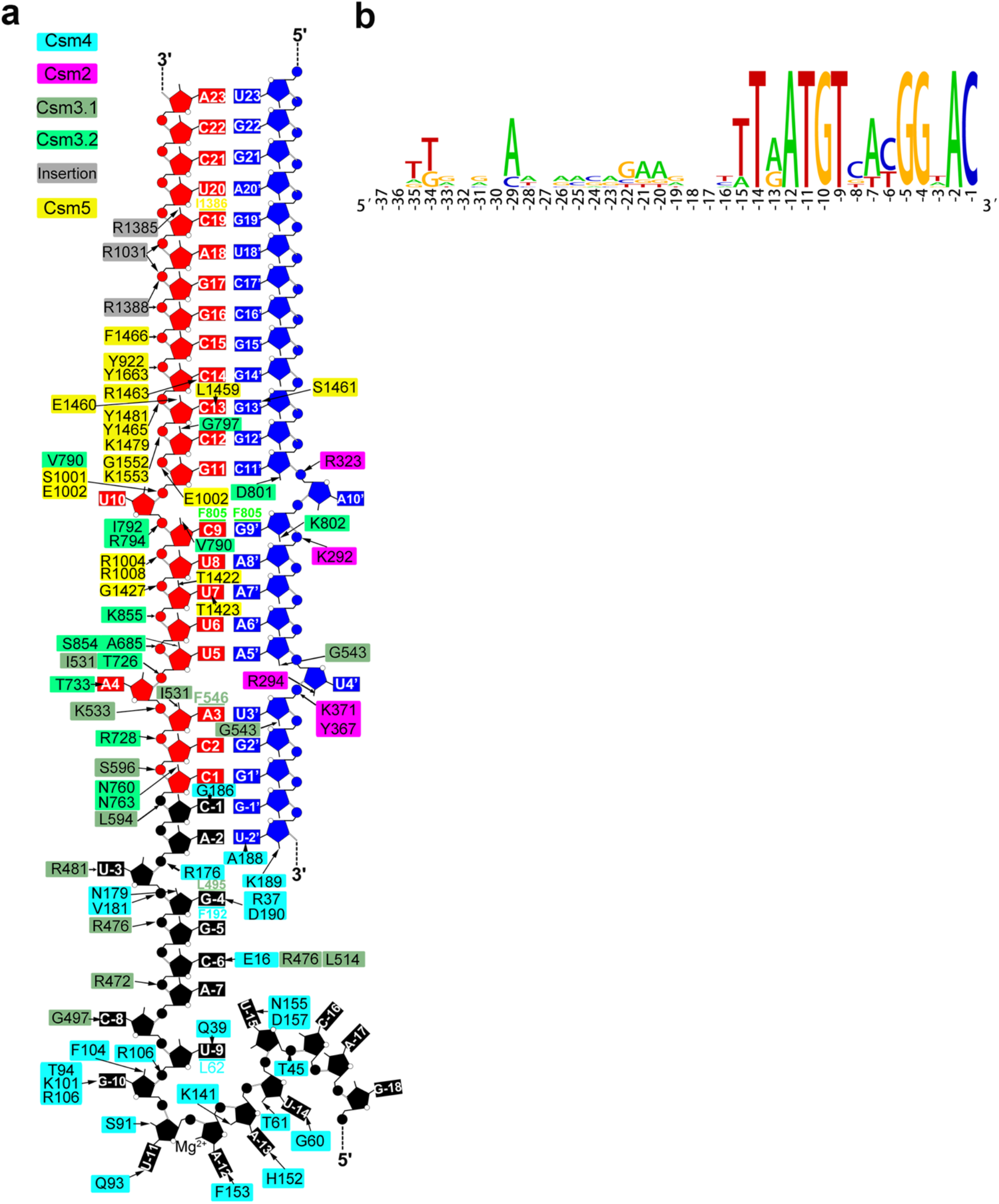
Recognition of crRNA-target RNA duplex by gRAMP. **a**, Detailed interactions between crRNA-target RNA duplex and gRAMP in gRAMP^crRNA^-target RNA^non-self^ ternary complex. **b**, Seqlogos of representative type III-E repeats constructed by WebLogo (http://weblogo.berkeley.edu/logo.cgi). Repeats are found by CRISPRminer (http://www.microbiome-bigdata.com/CRISPRminer) from MGTA01000040.1; PDWI01005922.1; SESD01000293.1; OBJA01001127.1; NZ_BEXT01000001.1; RHLA01000020.1; JRYO01000185.1.

**Extended Data Fig. 5.**
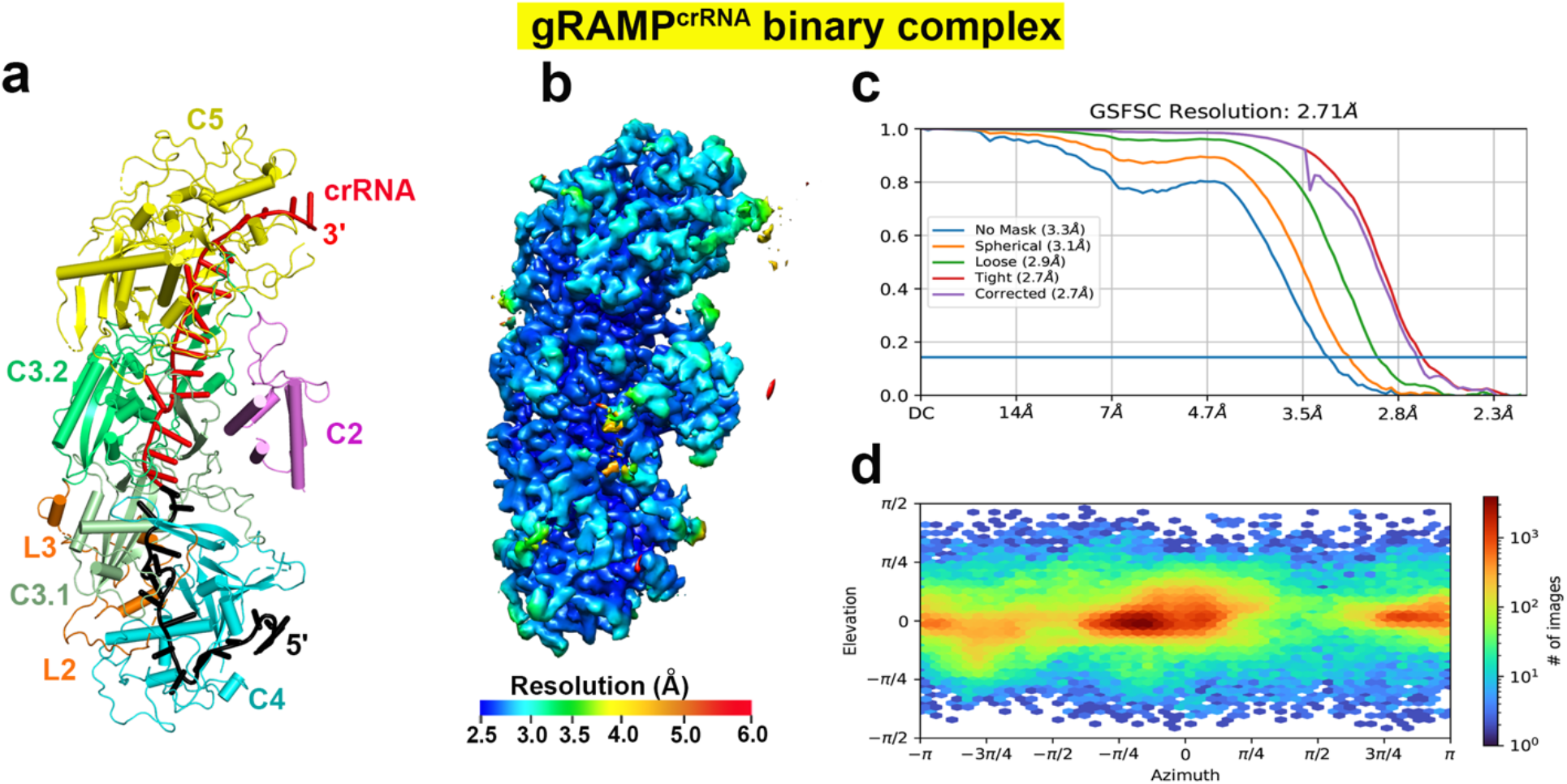
Cryo-EM Reconstruction of gRAMP^crRNA^ binary complex. **a**, Cryo-EM structure of gRAMP^crRNA^ binary complex. **b**, Final 3D reconstructed map of gRAMP^crRNA^-binary complex colored according to local resolution. **c**, FSC curve of gRAMP^crRNA^ binary complex reconstruction. **d**, Direction distribution plot of gRAMP^crRNA^ binary complex reconstruction.

**Extended Data Fig. 6.**
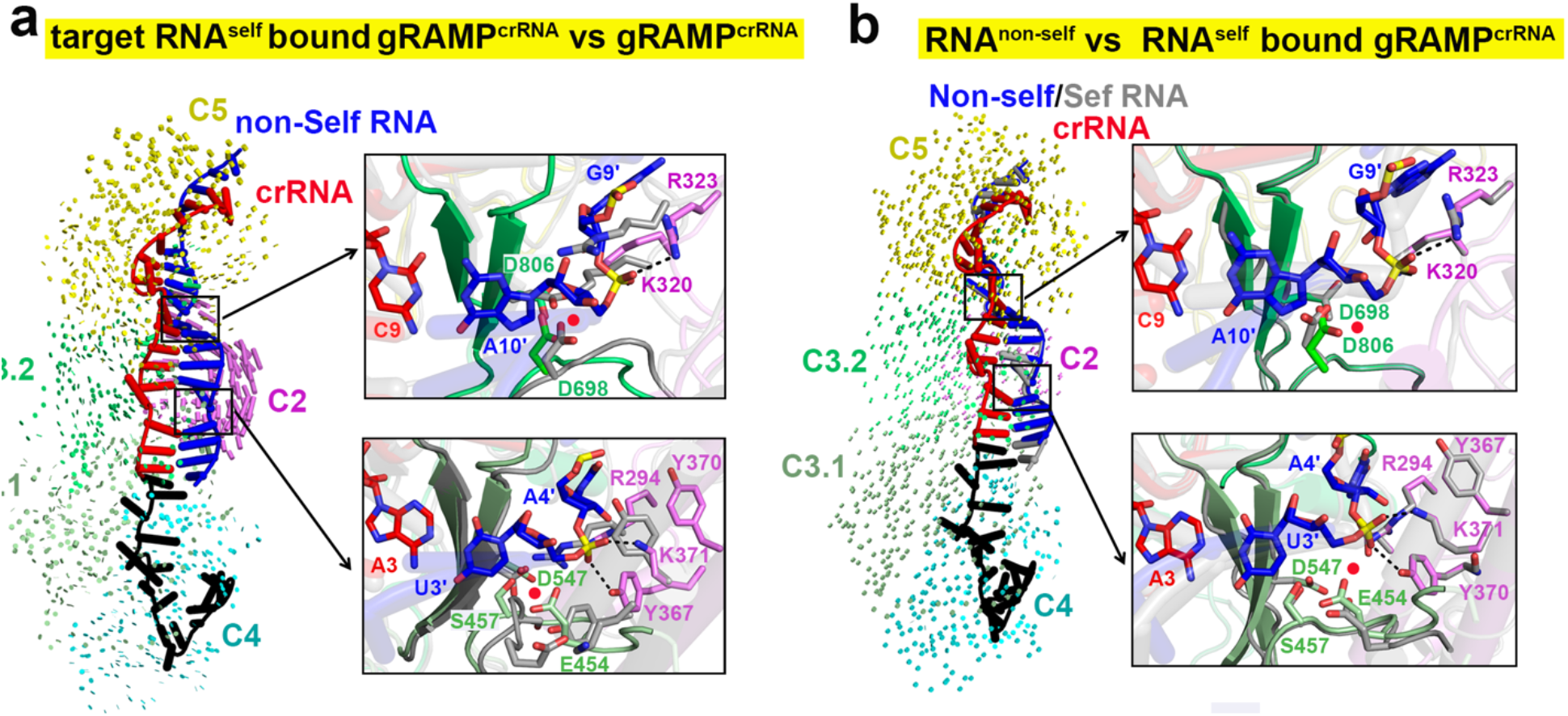
Structural comparison of RNase catalytic pockets in gRAMP complexes. **a**, Structure comparison of the overall structure and residues in the RNase catalytic pockets in gRAMP^crRNA^ binary (in grey) and gRAMP^crRNA^-target RNA^non-self^ ternary complex (in color). **b**, Structure comparison of the overall structure and residues in the RNase catalytic pockets in gRAMP^crRNA^-target RNA^self^ (in grey) and gRAMP^crRNA^-target RNA^non-self^ ternary cnon-complexes (in color).

**Extended Data Fig. 7.**
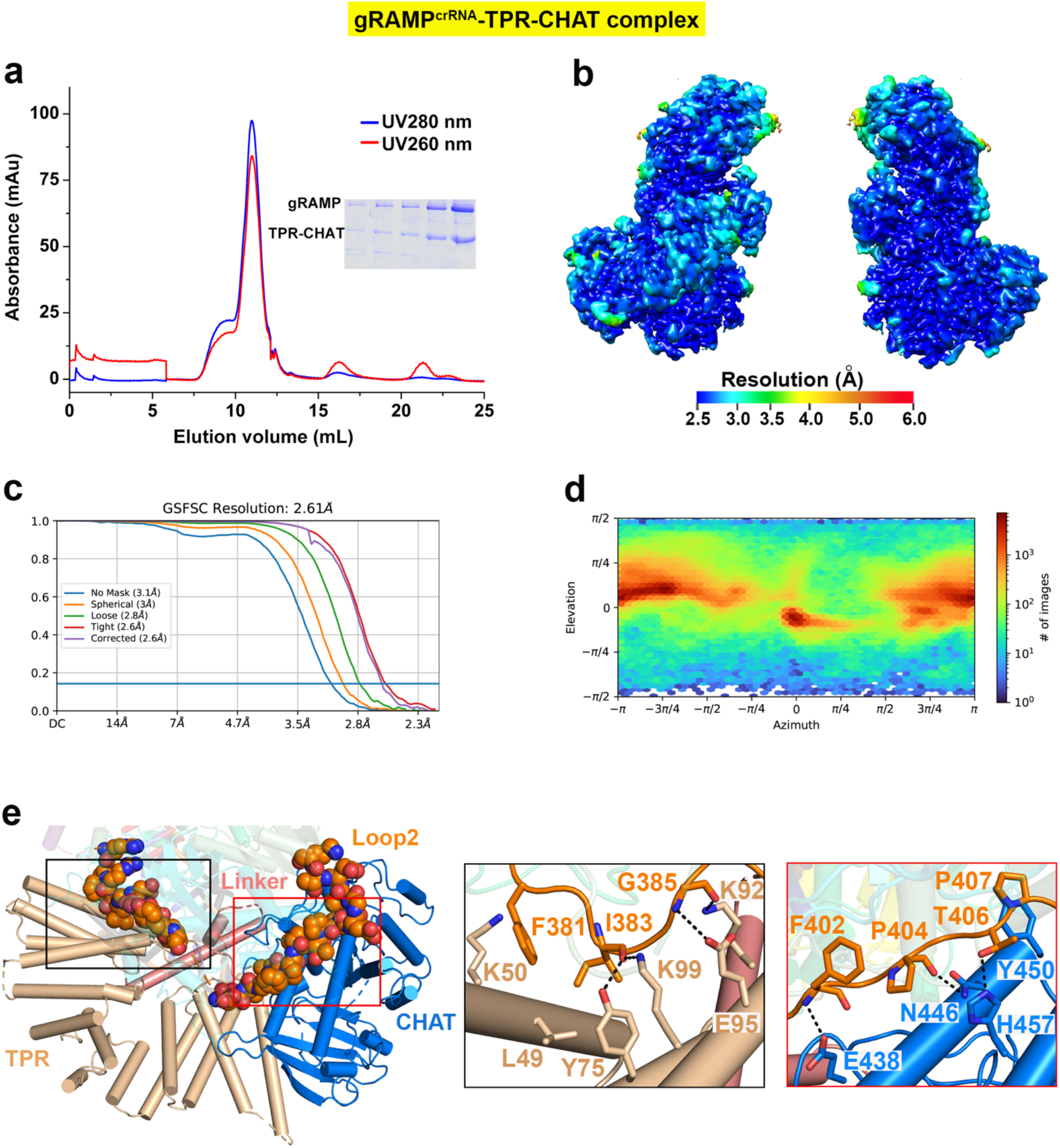
Reconstruction of gRAMP^crRNA^-TPR-CHAT complex. **a**, Size-exclusion chromatography and SDS-PAGE profile for purification of gRAMP^crRNA^-TPR-CHAT complex. Red and blue curves correspond to 260 and 280 nm UV absorptions, respectively. **b**, Final 3D reconstructed map of gRAMP^crRNA^-TPR-CHAT complex colored according to local resolution. **c**, FSC curve of gRAMP^crRNA^-TPR-CHAT complex reconstruction. **d**, Direction distribution plot of gRAMP^crRNA^-TPR-CHAT complex reconstruction. **e**, The detailed interactions between TPR-CHAT and Loop 2 in gRAMP in gRAMP^crRNA^-TPR-CHAT complex.

**Extended Data Fig. 8.**
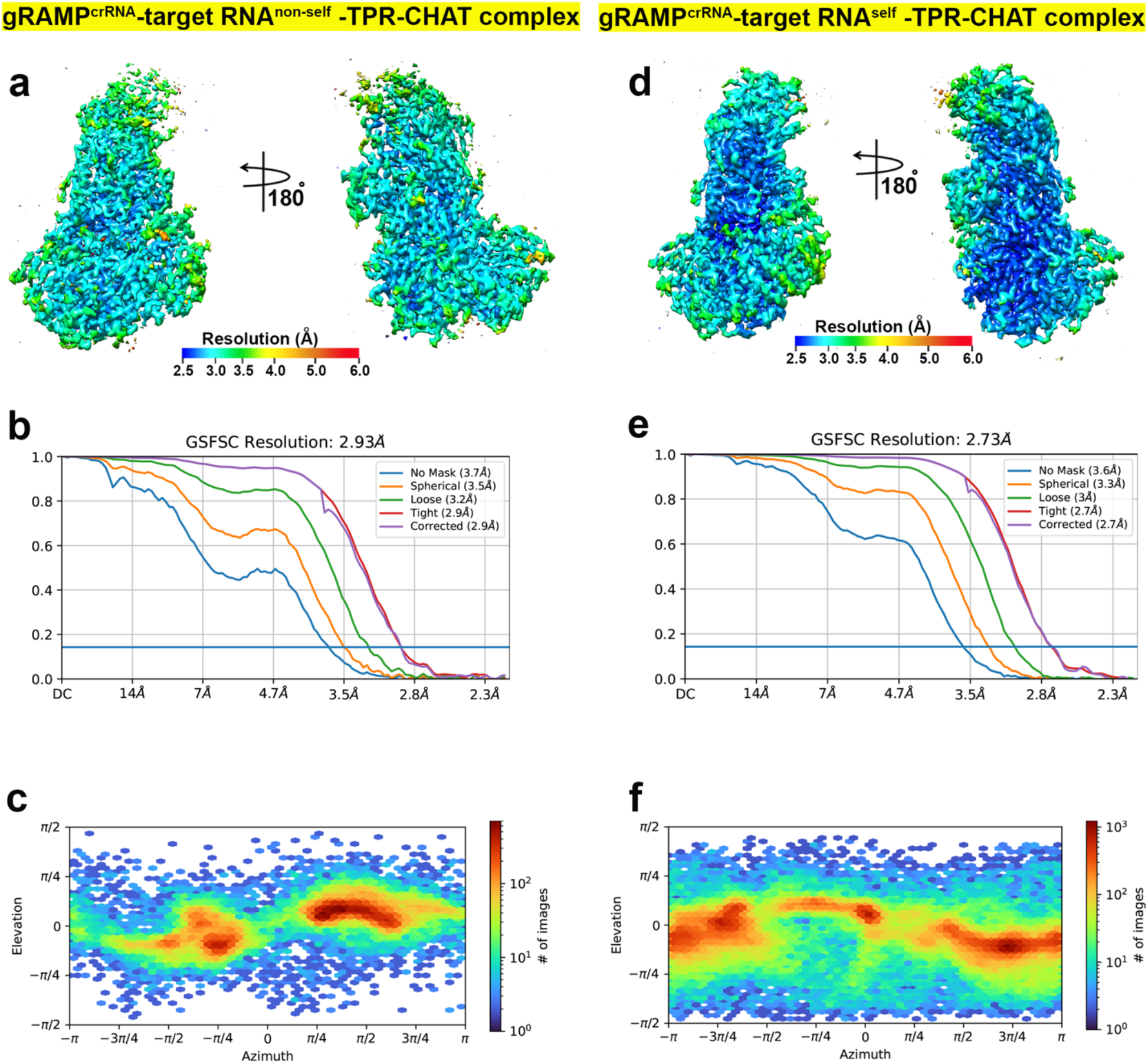
Cryo-EM Reconstruction of target RNA bound gRAMP^crRNA^-TPR-CHAT complex. **a** and **d**, Final 3D reconstructed map of gRAMP^crRNA^-TPR-CHAT-target RNA^non-self^ complex (**a**) and gRAMP^crRNA^-TPR-CHAT-target RNA^self^ complex (**d**) colored according to local resolution. **b** and **e**, FSC curve of gRAMP^crRNA^-TPR-CHAT-target RNA^non-self^ complex (**b**) and gRAMP^crRNA^-TPR-CHAT-target RNA^self^ complex (**e**) reconstruction. **c** and **f**, Direction distribution plot of gRAMP^crRNA^-TPR-CHAT-target RNA^non-self^ (**c**) and gRAMP^crRNA^-TPR-CHAT-target RNA^self^ (**f**) complex reconstruction.

**Extended Data Fig. 9.**
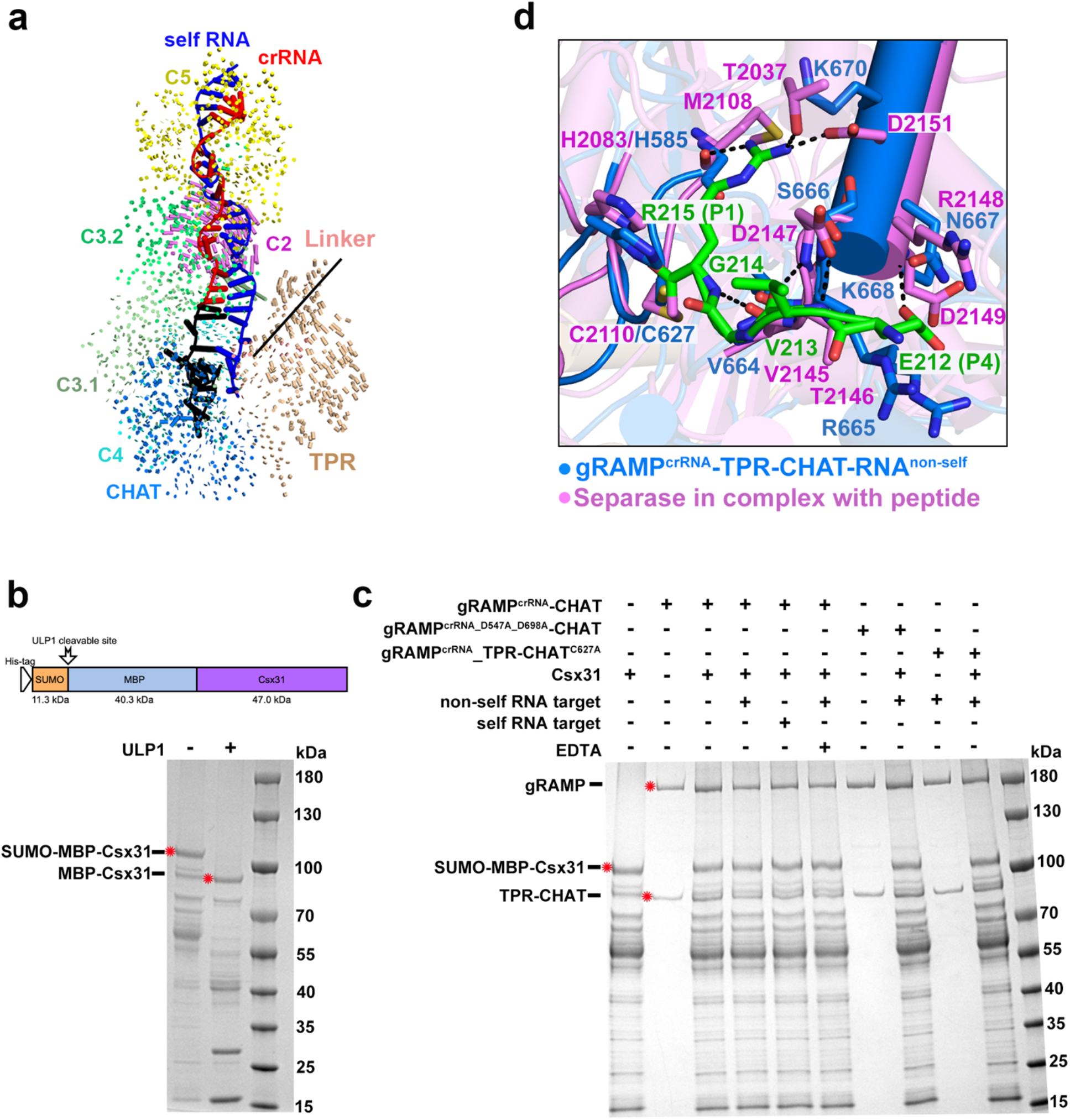
The protease activity of gRAMP^crRNA^-TPR-CHAT complex. **a**, Structure comparison between gRAMP^crRNA^-TPR-CHAT before and after self RNA target binding. Vector length indicates the domain movement scale. **b**, Cleavage of Csx31 fused with ULP1 Maltose binding protein (MBP) and SUMO tags demonstrated the expression and position of SUMO-MPB-Csx31 protein. We could only get the soluble Csx31 fused with SUMO and MPB tags. **c**, Cleavage of SUMO-MPB-Csx31 protein by the gRMAP^crRNA^-CHAT complex and its mutants. The red asterisk indicates the position of proteins labeled in left panel. **d**, Superposition of residues in the catalytic pockets between TPR-CHAT (in marine) and *Chaetomium thermophilum* separase (in violet, PDB 5FC3).

**Extended Data Table 1.** Cryo-EM data collection, refinement and validation statistics.

|  | gRAMP<sup>crRNA</sup> | gRAMP<sup>crRNA</sup><sub>target RNA<sup>non-self</sup></sub> | gRAMP<sup>crRNA</sup><sub>target RNA<sup>self</sup></sub> | gRAMP<sup>crRNA</sup><sub>TPR-CHAT</sub> | gRAMP<sup>crRNA</sup><sub>TPR-CHAT-target RNA<sup>non-self</sup></sub> | gRAMP<sup>crRNA</sup><sub>TPR-CHAT-target RNA<sup>self</sup></sub> |
| --- | --- | --- | --- | --- | --- | --- |
| **Data collection and processing** |  |  |  |  |  |  |
| Magnification | 105,000 | 105,000 | 105,000 | 105,000 | 105,000 | 105,000 |
| Voltage (kV) | 300 | 300 | 300 | 300 | 300 | 300 |
| Electron exposure (e<sup>-</sup>/Å<sup>2</sup>) | 50 | 50 | 50 | 50 | 50 | 50 |
| Defocus range (μm) | 1.5 to 2.5 | 1.5 to 2.5 | 1.5 to 2.5 | 1.5 to 2.5 | 1.5 to 2.5 | 1.5 to 2.5 |
| Pixel size (Å) | 1.1 | 1.1 | 1.1 | 1.1 | 1.1 | 1.1 |
| Symmetry imposed | C1 | C1 | C1 | C1 | C1 | C1 |
| Initial particle images (no.) | 2,895,967 | 3,310,582 | 1,610,246 | 3,120,917 | 1,792,636 | 2,002,116 |
| Final particle images (no.) | 227,997 | 444,936 | 76,820 | 611,350 | 53,439 | 123,862 |
| Map resolution (Å) | 2.71 | 2.54 | 2.83 | 2.61 | 2.93 | 2.73 |
| FSC threshold | 0.143 | 0.143 | 0.143 | 0.143 | 0.143 | 0.143 |
| Map resolution range (Å) | 2.2~999 | 2.2~999 | 2.2~999 | 2.2~999 | 2.2~999 | 2.2~999 |
| **Refinement** |  |  |  |  |  |  |
| Initial model used (PDB code) | *ab-initio* | *ab-initio* | *ab-initio* | *ab-initio* | *ab-initio* | *ab-initio* |
| Model resolution (Å) | 2.91 | 2.72 | 3.05 | 2.73 | 3.17 | 2.96 |
| FSC threshold | 0.5 | 0.5 | 0.5 | 0.5 | 0.5 | 0.5 |
| Map sharpening *B* factor (Å<sup>2</sup>) | -110.6 | -106.6 | -98.6 | -119.3 | -85.2 | -96.5 |
| Map Correlation Coefficient | 0.75 | 0.80 | 0.82 | 0.82 | 0.80 | 0.75 |
| **Model composition** |  |  |  |  |  |  |
| Non-hydrogen atoms | 10,572 | 11,044 | 12,278 | 16,081 | 16,292 | 16,502 |
| Protein residues | 1,218 | 1,223 | 1,341 | 1,894 | 1,860 | 1,884 |
| Nucleotide base | 35 | 55 | 66 | 35 | 59 | 59 |
| ***B* factor (Å<sup>2</sup>)** |  |  |  |  |  |  |
| Protein | 54.58 | 43.72 | 32.54 | 38.54 | 52.78 | 94.20 |
| Nucleotide | 51.10 | 47.35 | 45.25 | 40.12 | 63.31 | 93.04 |
| **R.m.s. deviations** |  |  |  |  |  |  |
| Bond lengths (Å) | 0.005 | 0.005 | 0.003 | 0.006 | 0.003 | 0.004 |
| Bond angles (°) | 0.883 | 0.784 | 0.576 | 0.644 | 0.571 | 0.572 |
| **Validation** |  |  |  |  |  |  |
| MolProbity score | 1.89 | 1.72 | 2.19 | 1.91 | 2.50 | 2.21 |
| Clash score | 8.42 | 6.73 | 15.97 | 14.68 | 17.67 | 15.48 |
| Poor rotamers (%) | 3.77 | 2.91 | 4.34 | 2.04 | 5.46 | 3.93 |
| **Ramachandran plot** |  |  |  |  |  |  |
| Favored (%) | 98.24 | 98.75 | 98.25 | 98.06 | 96.87 | 97.73 |
| Allowed (%) | 1.76 | 1.25 | 1.75 | 1.94 | 3.13 | 2.27 |
| Disallowed (%) | 0 | 0 | 0 | 0 | 0 | 0 |
| PDB | 7Y80 | 7Y81 | 7Y82 | 7Y84 | 7Y83 | 7Y85 |
| EMDB | 33676 | 33677 | 33678 | 33680 | 33679 | 33681 |

